# Divergent Effects of *APOE3* and *APOE4* Human Astrocytes on Key Alzheimer’s Disease Hallmarks in Chimeric Mice

**DOI:** 10.1101/2025.01.28.635271

**Authors:** Joan Cruz-Sese, Marta Mirón-Alcala, Maria Alfonso-Triguero, Jon Olalde, Leire Ruiz, Nuria Galbis-Gramage, Lorea Cortes, Laura Escobar, Pranav Preman, An Snellinx, Takashi Saito, Takaomi C Saido, Laura Saiz-Aúz, Alberto Rábano-Gutiérrez, Julia TCW, Alison Goate, Bart De Strooper, Elena Alberdi, Amaia M Arranz

## Abstract

Despite strong evidence supporting that both astrocytes and apolipoprotein E (APOE) play crucial roles in the pathogenesis and progression of Alzheimer’s disease (AD), the impact of astrocytes carrying different *APOE* variants on key AD pathological hallmarks remains largely unknown. To explore such effects in a human relevant context, we generated a chimeric model of AD. We transplanted isogenic *APOE3* or *APOE4* human induced pluripotent stem cell (hiPSC)-derived astrocyte progenitors into neonatal brains of AD model mice. We show that at five to six months after transplantation, transplanted cells have differentiated into mature astrocytes (h-astrocytes) that often integrate in upper layers of one cortical hemisphere. *APOE3* and *APOE4* h-astrocytes differentially express and secrete the APOE protein, which binds to Aβ plaques with an isoform-dependent affinity. Remarkably, *APOE3* h-astrocytes ameliorate Aβ pathology, Tau pathology and neuritic dystrophy. In contrast, *APOE4* h-astrocytes aggravate these AD processes. Moreover, *APOE3* and *APOE4* h-astrocytes modulate microglia responses to Aβ pathology in opposite ways. *APOE4* h-astrocytes enhance microglia clustering around Aβ plaques and exacerbate DAM state whereas *APOE3* h-astrocytes reduce microglia clustering and induce a more homeostatic state on plaque-associated microglia. These findings highlight a critical contribution of h-astrocytes not only to Aβ pathology but also to other key AD hallmarks in chimeric mice. In addition, our findings reveal that h-astrocytes with different *APOE* variants and the different forms of APOE they secrete have a crucial role in AD progression.

## INTRODUCTION

The *APOE4* variant of the apolipoprotein E gene (*APOE*) is the strongest genetic risk factor for developing late-onset Alzheimer’s disease (LOAD). *APOE* encodes three common alleles (ε2, ε3, ε4) characterized by varying risks for the development of AD ^1^. *APOE* ε3 is the most common and considered the neutral allele whereas *APOE* ε4 significantly increases AD risk in a gene dose-dependent manner and, conversely, *APOE* ε*2* is associated with lower risk for AD and later age at onset ^2^. Within the brain, APOE is predominantly expressed and produced by astrocytes in healthy conditions, and microglia and neurons also express and produce it when damage occurs ^3^. Although its primary function is to reduce intracellular cholesterol levels by effluxing lipids, APOE also plays integral roles in the overall brain health and in the development and progression of AD ^1,2^. Still how different *APOE* variants affect AD pathogenesis and progression and their effects on different brain cell types are largely unknown.

Recent studies are starting to reveal that *APOE* isoforms influence astrocyte and microglia molecular states and functions in the context of AD ^4–6^. Besides, in the case of astrocytes, various studies have shown that astrocytes with different *APOE* variants differentially affect AD associated features such as Aβ burden, Tau tangle pathology and neurodegeneration ^7–13^. Most of this knowledge comes from studies in either AD mouse models or human induced pluripotent stem cell (hiPSC)-derived *in vitro* models. While we have learned a lot from mouse models, prominent differences exist between rodent and human brain cells. In the case of astrocytes, human astrocytes (h-astrocytes) are 2-3 times larger and more complex, have more processes and encompass a higher number of synapses ^14–16^. At molecular level, human and mouse astrocytes display different gene expression profiles ^16^, and importantly, the *APOE* polymorphism does not exist in rodents. At functional level, h-astrocytes show different responses when exposed to inflammatory stimuli ^17–19^ and propagate calcium waves faster than rodent ones ^15,16,20^. Moreover, AD mouse models are unable to recapitulate some key human-specific cellular hallmarks of LOAD ^21^. Due to these critical species-specific differences, and in order to study human-specific aspects of AD, many groups are using the stem cell technology and hiPSCs from AD patients and healthy individuals. The ability to generate hiPSC-derived astrocytes ^22–24^ and other CNS cells is providing exciting opportunities to investigate their phenotypes and functions *in vitro* in the context of AD ^25^. Yet, hiPSC-derived neurons and glia grown in culture have limited maturation and lack essential components present in the brain which can lead to altered phenotypes and gene expression profiles significantly different from those in the human brain ^16,22,26^.

To address these challenges and to assess how *APOE3* and *APOE4* h-astrocytes affect main AD hallmarks within the live brain, we used a xenotransplantation model in which we engrafted hiPSC-derived astrocyte progenitors into the brain of AD model mice. In this chimeric model, we found that *APOE3* and *APOE4* h-astrocyte progenitors similarly differentiate into mature astrocytes and integrate mainly in upper layers of one cortical hemisphere. Interestingly, in AD chimeric mice, *APOE3* and *APOE4* h-astrocytes exhibit distinct patterns of APOE expression and secretion and affect major AD processes in divergent ways. In cortical regions with *APOE4* h-astrocytes there is increased Aβ pathology and Tau pathology, and enhanced plaque-associated microglia responses and neuritic dystrophy. In contrast, cortical regions hosting isogenic *APOE3* h-astrocytes show a decrease in these pathological hallmarks. Together, our *in vivo* results provide crucial evidence supporting that h-astrocytes harboring either *APOE3* or *APOE4* variants and the different forms of APOE they secrete drive improvements or aggravate not only Aβ pathology, but also other key features of AD such as Tau pathology and neuritic dystrophy. Moreover, they modulate microglia responses in divergent ways, having a more impactful role on AD progression and neurodegeneration than previously thought. In addition, our data validate the use of chimeric mice as a powerful technology to explore the contribution of h-astrocytes to AD and to investigate interactions of h-astrocytes with other brain cells.

## RESULTS

### Transplanted hiPSC-derived astrocyte progenitors integrate efficiently within the mouse host brain and differentiate to human astrocytes

To generate chimeric mice with human astrocytes, we used a similar approach as we used before ^5^. Briefly, we differentiated eight hiPSC lines from three AD patients carrying either the *APOE E4/E4 (APOE4)* alleles, or the corresponding corrected *APOE E3/E3 (APOE3)* isogenic lines (Table 1), into astrocyte progenitor cells (hAPCs) *in vitro* following an established protocol ^23^ (Fig. 1A). After 44 days in culture, we transplanted the td-Tomato expressing hAPCs into the brain of newborn wildtype and AD mice (Fig. 1A). A total of four groups of mice were generated: wildtype or AD mice transplanted with either *APOE3* or *APOE4* hAPCs.

**Figure 1.**
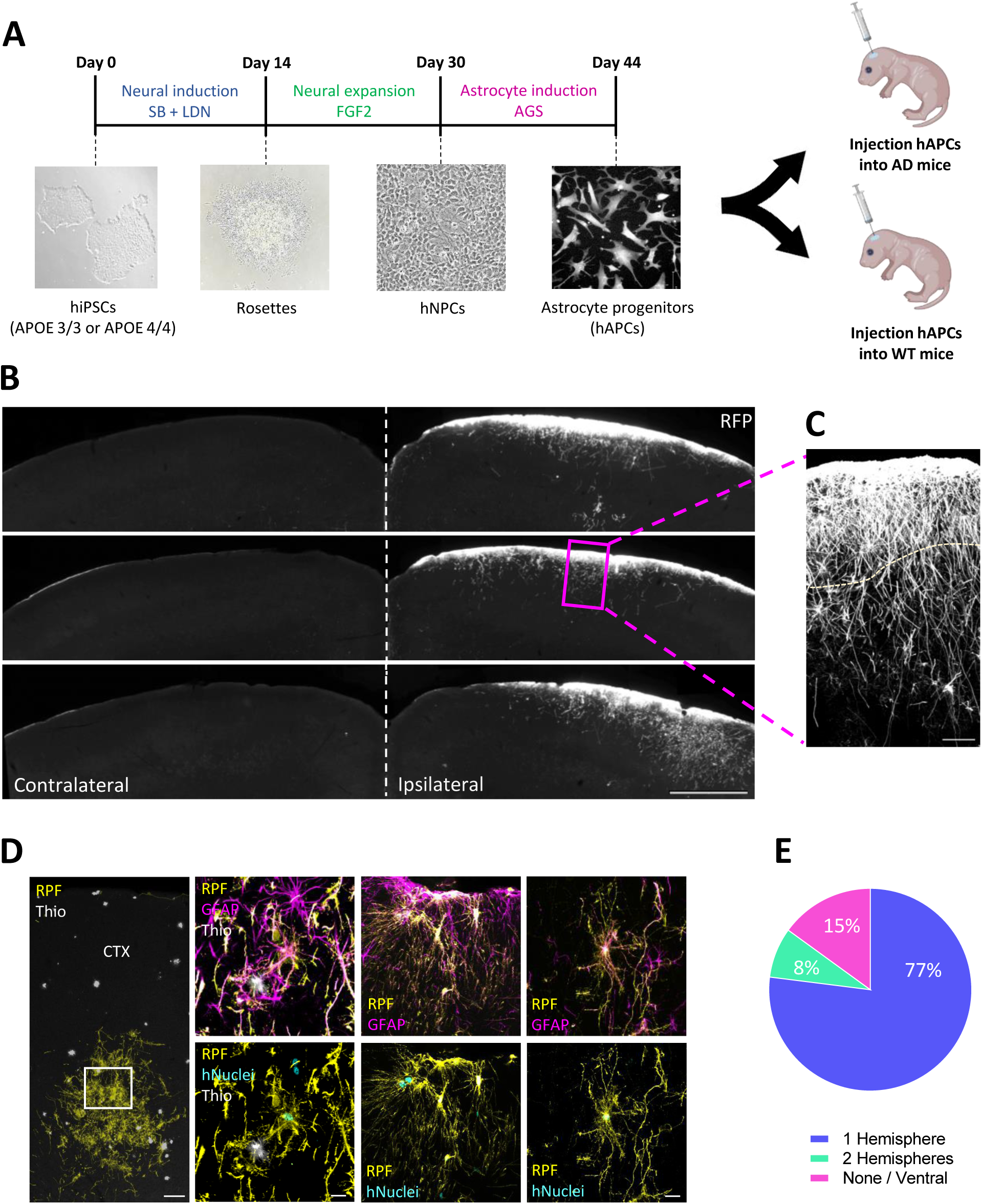
Transplanted cells integrate efficiently within the mouse host brain and differentiate to human astrocytes. **(A)** Schematics of the differentiation and transplantation procedures. hiPSCs: human induced pluripotent stem cells, hNPCs: human neural progenitor cells, hAPCs: human astrocyte progenitor cells, SB: SB431542, LDN: LDN193189, FGF2: fibroblast growth factor 2, AGS: astrocyte growth supplement. **(B)** Overview of the chimeric cortex showing h-astrocytes (RFP, white) in Layer I of one cortical hemisphere of chimeric brains 5 months after transplantation. Scale bar: 500 μm **(C)** High magnification view of h-astrocytes in mouse cortex Layer I. Scale bar: 50 μm. **(D)** Overview and higher magnification images of h-astrocytes (RFP, yellow; GFAP, magenta or hNuclei, cyan) integrated throughout the forebrain. Aβ plaques are stained with Thioflavin S (Thio, white). Scale bars: 20 μm in high magnification images and 100 μm in overview. **(E)** Pie chart showing the percentage of chimeric brains with cortical integration of h-astrocytes in one hemisphere, in the two hemispheres or no cortical integration (n= 53 mice).

**Table 1:** Information of the hiPSC lines.

| hiPSC name | Ethnicity | Sex | Age of onset | Age at skin biopsy | Disease status (CDR at biopsy) | *APOE* genotype | Genetic modification |
| --- | --- | --- | --- | --- | --- | --- | --- |
| TCW1E33-1F1 | Caucasian | F | 64 | 72 | AD (2) | E4/E4 | E3/E3 |
| TCW1E44-2C2 | Caucasian | F | 64 | 72 | AD (2) | E4/E4 | E4/E4 |
| TCW2E33-2E3 | Caucasian | M | 77 | 80 | AD (0.5) | E4/E4 | E3/E3 |
| TCW2E44-4B1 | Caucasian | M | 77 | 80 | AD (0.5) | E4/E4 | E4/E4 |
| TCW2E33-3D11 | Caucasian | M | 77 | 80 | AD (0.5) | E4/E4 | E3/E3 |
| TCW2E44-4B12 | Caucasian | M | 77 | 80 | AD (0.5) | E4/E4 | E4/E4 |
| TCW3E33-H-2 | Caucasian | M | 80 | 83 | AD (0.5) | E4/E4 | E3/E3 |
| TCW3E44-F-2 | Caucasian | M | 80 | 83 | AD (0.5) | E4/E4 | E4/E4 |

The table shows hiPSC name, patient ethnicity, sex, age of AD onset, age at skin biopsy, disease status (CDR at biopsy), original *APOE* genotype and genetic modification. F female, M male, AD Alzheimer's disease, *APOE* apolipoprotein, CDR clinical dementia rating, hiPSC human induced pluripotent stem cells. These cells were previously generated and characterized by TCW et al, 2022 <sup>11</sup> and Preman et al, 2021 <sup>5</sup>.

The chimeric mice were aged for five to six months and their brains were collected for immunofluorescence (IF) or protein extraction and MSD-ELISA analyses. Transplanted cells were identified through IF analyses based on the expression of the td-Tomato marker RFP and of the human nuclear antigen hNuclei (Fig S1A); they integrated efficiently within the mouse host brain and were able to contact blood vessels (Fig S1B) as we described before ^5^. Both *APOE3* and *APOE4* cells followed a similar pattern of engraftment with transplanted cells infiltrating throughout the forebrain (Fig 1D). In nearly 80% of the chimeric mice, a large proportion of the transplanted cells integrated in upper layers of one cortical hemisphere (Fig 1B, 1E) and showed astrocyte identity: they expressed main markers of astrocytes such as GFAP, vimentin, aquaporin and S100b (Fig S1C) and showed no expression of markers of neurons or oligodendrocytes (Fig S1D). Notably, they showed morphological features of interlaminar astrocytes present in Layer 1 of the human cortex ^27,28^, with small and round cell bodies near the pial surface (Fig S1A) and long and unbranched processes descending into deeper cortical layers (Fig 1B-C, S1A-D) in line with our previous published studies in chimeric mice ^5^. Together, these data indicate successful integration of transplanted hAPCs and maturation into human astrocytes (h-astrocytes) in both wildtype and AD mouse cortex. For all following analyses, we used the nearly 80% of chimeric brains with h-astrocytes integrated in upper layers of one cortical hemisphere (Fig 1B, 1E).

### Human astrocytes express and secrete APOE in an isoform-dependent manner

The APOE protein is mainly expressed and produced by astrocytes in homeostatic conditions with both microglia and neurons also expressing APOE in pathological conditions^29^. Therefore, we wondered whether h-astrocytes were expressing human APOE (hAPOE) *in vivo*, in the chimeric mice, and if there were isoform-dependent differences in hAPOE expression. We analyzed the expression of hAPOE within the h-astrocyte transplants by IF using a monoclonal antibody that specifically detects human APOE. We found an overall increase in hAPOE expression in *APOE3* h-astrocytes compared to *APOE4* h-astrocytes in both wildtype and AD mice (Fig 2A, C), in agreement with previous studies in h-astrocytes in culture ^10^. Intensity profiles of hAPOE IF signals across h-astrocytes confirmed these results and showed that while hAPOE signals were homogeneous across *APOE3* h-astrocytes, they were irregular in *APOE4* h-astrocytes with few cells highly expressing hAPOE and many cells showing very low expression (Fig S2A, B). Moreover, we found that while *APOE3* h-astrocytes significantly upregulated hAPOE in AD mice compared with the wildtype situation, hAPOE expression was not significantly modified in *APOE4* h-astrocytes in AD compared with WT mice (Fig 2A, C).

**Figure 2.**
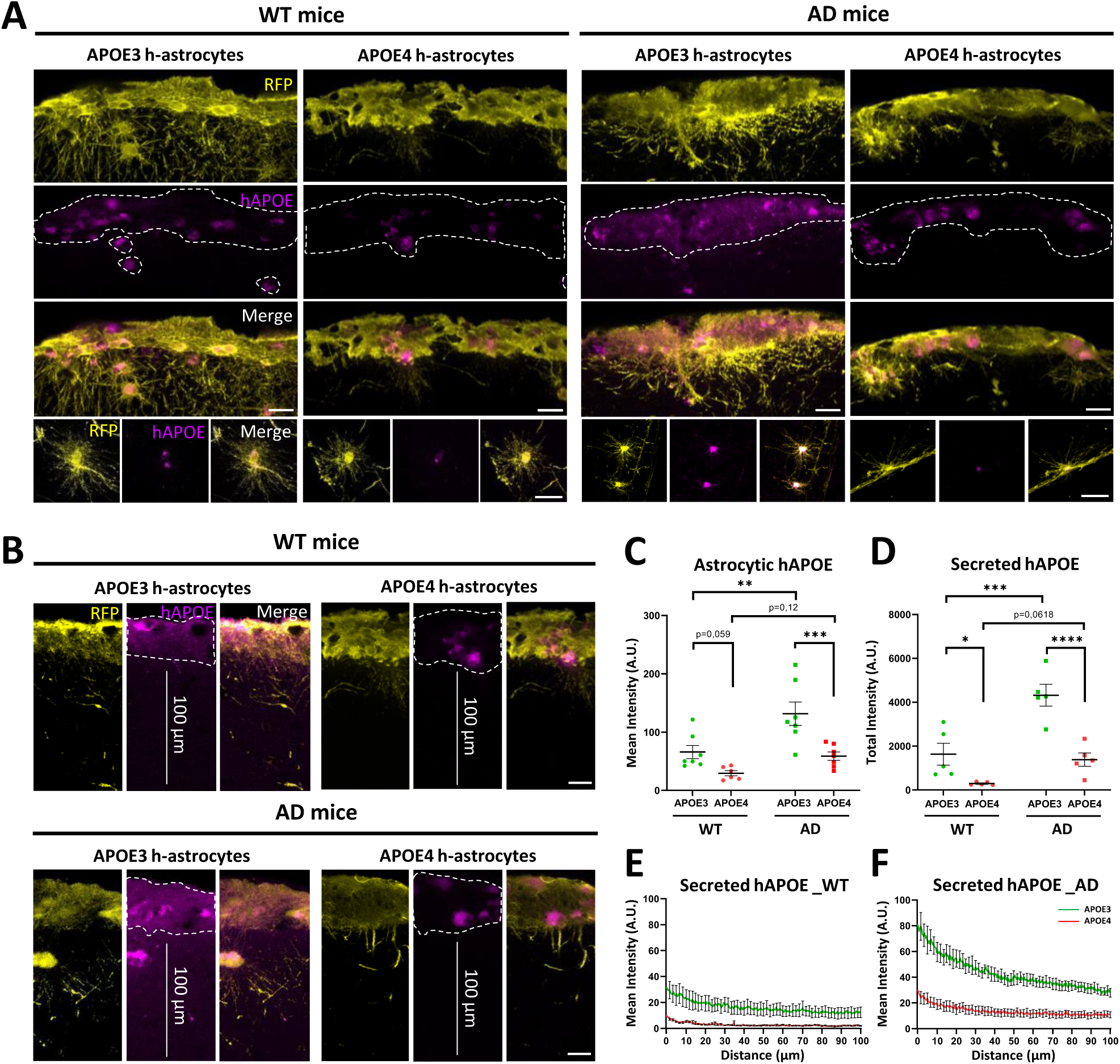
Human astrocytes express and secrete APOE in an isoform-dependent manner. (A, **B)** Representative images of the chimeric cortex showing *APOE3* and *APOE4* h-astrocytes (RFP, yellow) and human APOE (hAPOE, magenta) in 5-6 month old WT and AD mice. Scale bars: 20 μm in top panels and 200 μm in bottom panels. **(C)** Quantification of the hAPOE intensity per area in *APOE3* and *APOE4* h-astrocytes in WT and AD mice (n=7 WT mice with *APOE3* and n=6 WT mice with *APOE4* h-astrocytes; n=7 AD mice with *APOE3* and n=7 AD mice with *APOE4* h-astrocytes). **(D)** Quantification of the total intensity of secreted hAPOE in WT and AD mice hosting *APOE3* and *APOE4* h-astrocytes (n=5 WT mice with *APOE3* and n=5 WT mice with *APOE4* h-astrocytes; n=5 AD mice with *APOE3* and n=5 AD mice with *APOE4* h-astrocytes). Data are represented as mean ± SEM, 2-way ANOVA with Holm-Sidak test; ns = not significant, * p < 0.05, ** p < 0.01, *** p < 0.001. **(E, F)** Graphs representing the mean intensity of secreted hAPOE along a distance of 100 µm from the soma of h-astrocytes in both WT and AD mice.

Since APOE is secreted to the extracellular space, we next examined whether there were isoform-dependent differences in hAPOE secreted by h-astrocytes. We measured by IF the expression of secreted hAPOE in an area of 100 µm from the h-astrocyte transplants and found increased levels of secreted hAPOE in regions close to *APOE3* h-astrocytes compared to regions next to *APOE4* h-astrocytes (Fig 2B, D). Besides, levels of secreted hAPOE were upregulated in areas close to *APOE3* h-astrocytes in AD mice compared with wildtype mice, but were not significantly changed close to *APOE4* h-astrocytes in AD vs wildtype mice (Fig 2B, D-F). To confirm these data, we quantified hAPOE protein levels in homogenized cortex samples from chimeras using a human APOE specific ELISA and found that total hAPOE levels were significantly increased in cortex samples with *APOE3* h-astrocytes compared with samples hosting *APOE4* h-astrocytes (Fig S2C).

Overall, these data point to an isoform-dependent regulation of hAPOE expression and secretion by h-astrocytes *in vivo*, both in control and AD conditions, with reduced hAPOE expression and secretion in *APOE4* h-astrocytes compared with isogenic *APOE3* h-astrocytes.

### APOE from h-astrocytes show an isoform-dependent affinity binding to A**β** plaques

Interestingly, we noticed that hAPOE colocalized with amyloid-beta (Aβ) plaques in AD chimeric mice. We found hAPOE at the plaques in areas close to h-astrocytes (Fig 3A), both in Aβ plaques in direct contact with h-astrocyte processes and in plaques not contacting h-astrocytes directly (Fig 3B). This indicates that both hAPOE secreted by h-astrocytes to the extracellular space and hAPOE from h-astrocyte processes directly contacting the plaques bind to Aβ plaques (Fig 3B). However, we could not detect the presence of hAPOE in Aβ plaques far from h-astrocytes or in the contralateral cortical hemisphere devoid of h-astrocytes (Fig S2D). We measured the levels of hAPOE within plaques close to h-astrocyte transplants (in an area of 100 µm from the h-astrocytes) and remarkably found similar hAPOE levels in plaques close to *APOE3* and *APOE4* h-astrocytes (Fig 3C, D). These data reveal that *APOE3* h-astrocytes express and secrete more hAPOE than *APOE4* h-astrocytes and upregulate hAPOE expression and secretion in AD compared with wildtype chimeric mice. However, similar levels of hAPOE are found at the plaques, suggesting that hAPOE secreted by *APOE4* h-astrocytes might bind with higher affinity to Aβ plaques. In line with this, we quantified the ratio of hAPOE within the plaques versus secreted hAPOE surrounding the plaques and found a higher ratio in cortical areas close to *APOE4* h-astrocytes than in areas next to *APOE3* h-astrocytes (Fig 3C, E). These data confirm a higher affinity of hAPOE4 to bind and accumulate within the plaques.

**Figure 3.**
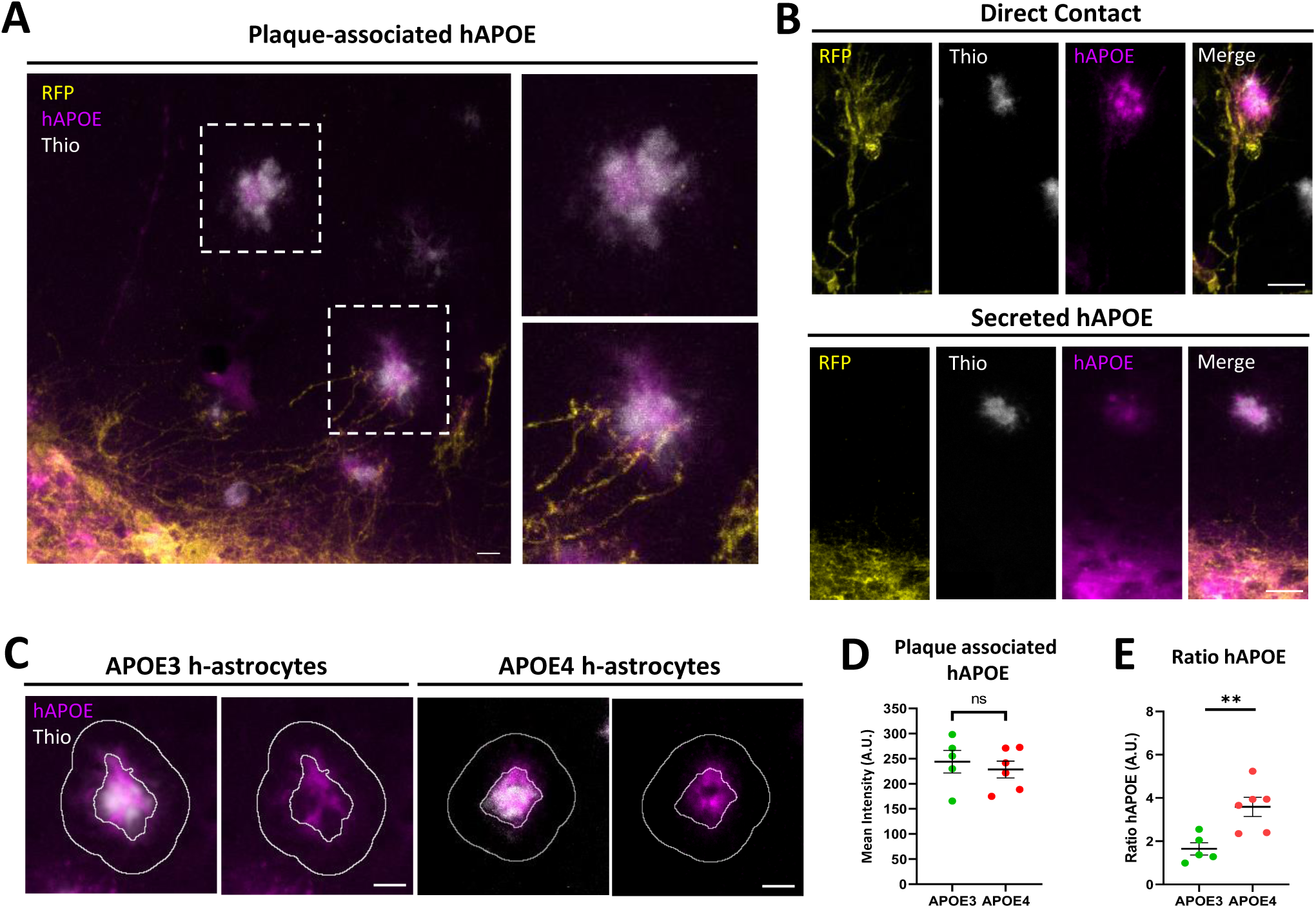
APOE from h-astrocytes binds to. **A**β **plaques with an isoform-dependent affinity. (A)** Representative images of h-astrocytes and their processes (RFP, yellow) and human APOE (hAPOE, magenta) that binds to Aβ plaques (Thio, white). Scale bars: 20 μm. **(B)** Both hAPOE from h-astrocyte processes in direct contact with the plaques (top panel) and hAPOE secreted to the extracellular space (bottom panel) binds to Aβ plaques. Scale bars: 20 μm. **(C)** Representative images of human APOE (hAPOE, magenta) within and surrounding Aβ plaques (Thio, white). Scale bars: 10 μm. **(D)** Quantification of plaque-associated hAPOE mean intensity in AD chimeras with *APOE3* and *APOE4* h-astrocytes (n=5 AD mice with *APOE3* and n=6 AD mice with *APOE4* h-astrocytes). **(E)** Ratio of hAPOE intensity within the plaques vs hAPOE intensity surrounding the plaques (n=5 AD mice with *APOE3* and n=6 AD mice with *APOE4* h-astrocytes). Data are represented as mean ± SEM, unpaired t-test; ns = not significant, * p < 0.05, ** p < 0.01, *** p < 0.001.

### *APOE3* and *APOE4* h-astrocytes differentially affect Aβ pathology

The way h-astrocytes integrate in the host brain, with most of the human cells in upper layers of one cortical hemisphere (Fig 1B, E), allowed analyzing their contribution to key AD pathological hallmarks within the same animal and brain section. Thus, we could compare the cortical hemisphere hosting h-astrocytes with the contralateral hemisphere that lacks transplanted cells (Fig 1B). We previously reported that transplanted h-astrocytes were able to closely interact with Aβ plaques in AD mouse brains ^5^. This provides a unique opportunity to study whether astrocytes with different *APOE* genotypes could differentially affect Aβ pathology. We used the dense core plaque marker Thioflavin S to stain the fibrillar Aβ plaques in cortical sections of 5-6 month-old chimeric AD mice. Intriguingly, we found a significant decrease in Aβ load in cortical regions hosting *APOE3* h-astrocytes compared with contralateral regions (Fig 4A, B). However, areas with *APOE4* h-astrocytes had more Aβ plaques per area compared with contralateral regions but did not show significant differences in the area covered by Aβ plaques or in the Aβ plaque size (Fig 4A, B). To assess overall Aβ deposition, we performed staining with the Aβ-specific monoclonal antibody 6C3. As with Thioflavin S, total amyloid burden decreased in cortical regions hosting *APOE3* h-astrocytes compared with contralateral regions (Fig 4C, D), while there were more Aβ plaques per area in regions where *APOE4* h-astrocytes were present compared with their contralateral sides (Fig 4C, D). We also quantified Aβ load in cortical areas close to h-astrocytes (within 100 µm from the transplanted cells) (Fig S3A), and found a similar trend (Fig S3A, B). These data reveal that not only the h-astrocytes or their processes but also the hAPOE they secrete influence Aβ pathology in the chimeric brains.

**Figure 4.**
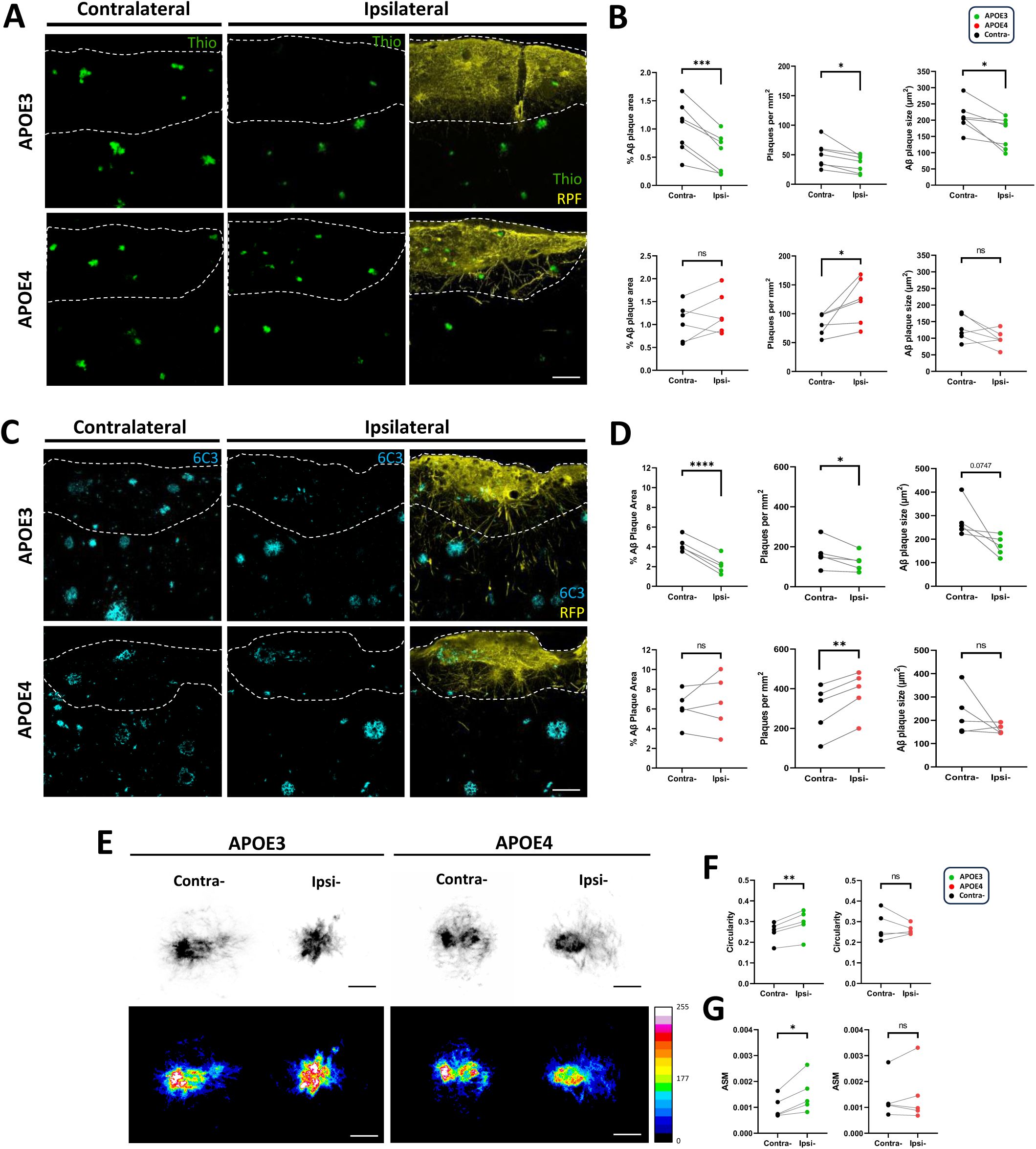
*APOE3* and *APOE4* h-astrocytes differentially affect. **A**β **pathology. (A)** Representative images of the cortex of AD chimeric mice 5-6 months after transplantation stained for h-astrocytes (RFP, yellow) and fibrillar Aβ (Thio, green). Both contralateral and ipsilateral cortices are shown. Scale bars: 50 μm. **(B)** Quantification of Aβ burden, number of plaques per area and plaque size in cortex regions hosting *APOE3* or *APOE4* h-astrocytes and in contralateral cortex (n=7 mice with *APOE3* and n=6 mice with *APOE4* astrocytes). **(C)** Representative images of the cortex of AD chimeric mice stained for h-astrocytes (RFP, yellow) and total Aβ (6C3, magenta). Scale bars: 50 μm. **(D)** Quantification of Aβ burden, number of plaques per area and plaque size in cortex regions with *APOE3* or *APOE4* h-astrocytes and in contralateral areas (n=5 mice with *APOE3* and n=5 mice with *APOE4* h-astrocytes). Data are represented as mean ± SEM, Paired t-test. **(E)** Representative images of X34-positive Aβ plaque conformation in grayscale and heat-density map based on intensity (red represents the highest intensity and blue is the lowest intensity) in contra- and ipsilateral cortices with *APOE3* and *APOE4* h-astrocytes. Scale bars: 10 μm**. (F, G)** Quantification of circularity (F) and ASM degree of compactness (G) of X34-positive plaques in contra- and ipsilateral cortices (n=5 mice with *APOE3* and n=5 mice with *APOE4* h-astrocytes). Data are represented as mean ± SEM, Paired t-test; ns = not significant, * = p < 0.05, ** = p < 0.01, *** = p < 0.001.

To further analyze the accumulation of Aβ in these cortical regions, we next used ELISA to quantify both soluble and insoluble Aβ_42_ levels. In AD chimeras, both soluble and insoluble Aβ_42_ levels significantly decreased in areas hosting *APOE3* h-astrocytes compared with contralateral areas (Fig S3C). However, we could not detect significant differences neither in soluble nor in insoluble Aβ_42_ levels in the presence of *APOE4* h-astrocytes compared with the contralateral side (Fig S3C). We also measured both soluble and insoluble Aβ_42_ levels in cortical regions hosting *APOE3* or *APOE4* h-astrocytes in wildtype chimeras. We found no or minimal levels of soluble and insoluble Aβ_42_ (Fig S3D), indicating that h-astrocytes, irrespective of their *APOE* genotype, were not producing Aβ and contributing to plaque burden in our model at this stage. Our data suggest that both *APOE3* and *APOE4* h-astrocytes and the hAPOE they secrete can affect the deposition of Aβ into fibrillar Aβ plaques. Therefore, we investigated potential changes in plaque morphology and conformation. We quantified circularity and compactness of individual Aβ plaques stained with X-34 in cortical areas near the h-astrocyte transplants. Interestingly, in Aβ plaques containing hAPOE3 and close to *APOE3* h-astrocytes, plaque circularity and compactness increased compared with plaques in contralateral areas (Fig 4E-G). However, there were no significant differences in plaque circularity and compactness in hAPOE4 containing plaques close to *APOE4* h-astrocytes compared with plaques in contralateral sides (Fig 4E-G).

Together, these results point to a critical contribution of h-astrocytes and secreted hAPOE to Aβ pathology *in vivo* in chimeric AD mice and reveal a distinct impact of h-astrocytes with different *APOE* variants. *APOE3* h-astrocytes and hAPOE3 significantly reduce plaque burden and increase plaque compactness. In contrast, *APOE4* h-astrocytes and hAPOE4 have a moderate impact on Aβ load increasing the number of plaques per area without affecting plaque size and compactness.

### Human brains from AD patients harboring *APOE4* alleles exhibit increased A**β** load and plaque-associated hAPOE ratio

To assess the translatability of our findings, we analyzed Aβ load and hAPOE expression at the plaques in the lateral temporal cortex from a cohort of either *APOE3/3 (APOE3)* or *APOE4/4 (APOE4)* carriers with a neuropathological diagnosis of AD at Braak stages V and VI (Table 3).

To analyze Aβ burden, we used Thioflavin S to stain the Aβ plaques. We observed that *APOE3* patients showed reduced Aβ load compared to *APOE4* patients in these cortical regions (Fig S4A, B), in agreement with previous studies in AD patients ^30^ and with our data in chimeric mice (Fig 4A,B). Next, we analyzed hAPOE expression at the plaques with the same human specific antibody, as well as the ratio of hAPOE at the plaques versus secreted hAPOE around the plaques. Interestingly, we detected hAPOE in Aβ plaques in AD patients (Fig S4A) and observed higher hAPOE ratios in *APOE4* than in *APOE3* patients (Fig S4D), as we found in chimeric mice. These data indicate a higher affinity of hAPOE4 to bind and accumulate within the plaques in *APOE4* patients. Additionally, we found higher hAPOE levels at the plaques in *APOE4* patients compared with *APOE3* patients (Fig S4C), in contrast with our results in chimeric mice, in which we found similar levels of hAPOE at the plaques (Fig 3D). These differences are probably due to the fact that in AD patients hAPOE is produced not only by astrocytes but also by microglia and neurons that upregulate hAPOE expression in AD ^3^.

### *APOE3* and *APOE4* h-astrocytes and secreted hAPOE differentially modulate microglial responses

Aβ pathology is accompanied by the clustering of microglia around amyloid plaques. Since microglial responses can be influenced by *APOE* isoforms ^9,31–33^, we next asked whether *APOE3* and *APOE4* h-astrocytes and the hAPOE they secrete could affect the responses of microglial cells to amyloid plaques in our chimeric mice. We performed IF with Iba-1, a myeloid cell marker upregulated in microglia, and Thioflavin S to stain the Aβ plaques. Then, we analyzed microglia clustering around each Aβ plaque in cortical regions within 100 µm from h-astrocytes and contralateral cortical areas. Microglial clustering was analyzed in Aβ plaques of similar sizes (Fig S7B) to rule out the possibility that changes observed in microglia were associated to plaque size. Interestingly, in cortical areas close to *APOE3* h-astrocytes, clustering of microglia around hAPOE positive plaques decreased compared with contralateral regions (Fig 5A, B). However, in areas next to *APOE4* h-astrocytes, microglia clustering around hAPOE containing plaques increased compared with the contralateral side (Fig 5A, B). We also used the microglial marker Iba-1 to examine total microglia, both plaque-associated and non-plaque-associated microglia. We found that while the area of the cortex covered by total microglia decreased in regions close to *APOE3* h-astrocytes, it remained unchanged in regions next to *APOE4* h-astrocytes compared with their respective contralateral sides (Fig S5A, B).

**Figure 5.**
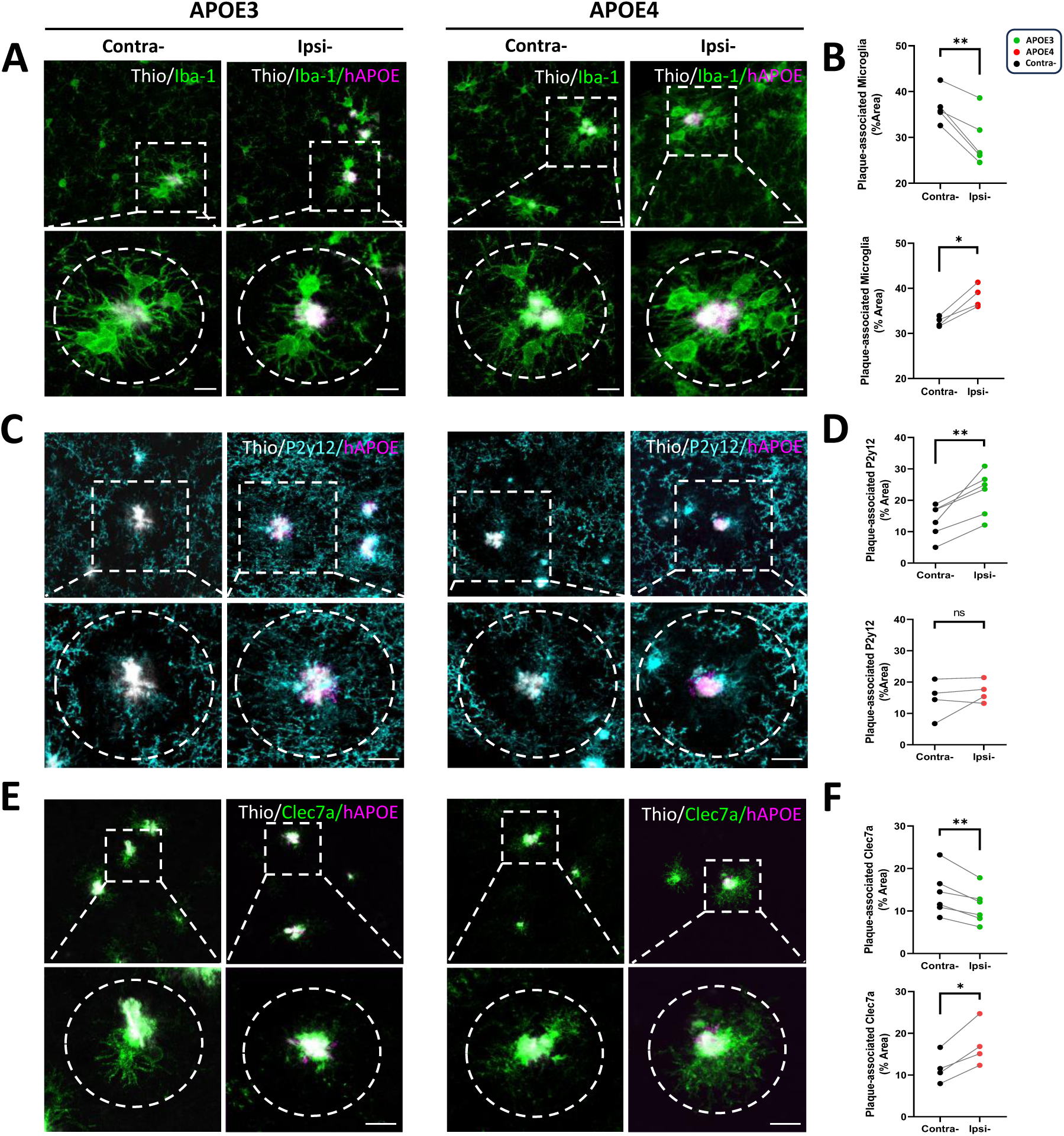
*APOE3* and *APOE4* h-astrocytes and secreted hAPOE differentially modulate microglial responses. **(A)** Representative images Iba-1 (green) positive microglia around Aβ plaques (Thio, white) with or without human APOE (hAPOE, magenta) in contralateral and ipsilateral cortices of AD chimeras hosting *APOE3* and *APOE4* h-astrocytes 5-6 months after transplantation. Scale bars: 10 μm. **(B)** Percentage area covered by Iba-1 positive microglia around Aβ plaques in AD chimeras (n=5 mice with *APOE3* and n=4 mice with *APOE4* h-astrocytes). **(C)** Representative images of P2y12 (cyan) positive microglial cells surrounding Aβ plaques (Thio, white) with and without human APOE (hAPOE, magenta) in contra- and ipsilateral cortices of AD chimeras. Scale bars: 10 μm. **(D)** Percentage area covered by P2y12 positive microglia around plaques (n=6 mice with *APOE3* and n=4 mice with *APOE4* h-astrocytes). **(E)** Representative images of Clec7a (green) positive microglial cells surrounding Aβ plaques (Thio, white) with and without human APOE (hAPOE, magenta) in contra- and ipsilateral cortices of AD chimeras. Scale bars: 10 μm. **(F)** Percentage area covered by Clec7a positive microglia around plaques (n=6 mice with *APOE3* and n=4 mice with *APOE4* h-astrocytes). Data are represented as mean ± SEM, paired t-test; ns = not significant, * = p < 0.05, ** = p < 0.001, *** = p < 0.0001.

These changes in microglia responses prompted us to investigate microglia activation around plaques. We therefore analyzed the expression of P2y12 and Clec7a, which are markers of homeostatic and disease-associated microglia (DAM) respectively. While cortical areas close to *APOE3* h-astrocytes showed a significantly higher P2y12 coverage of the plaques compared with contralateral regions, plaque coverage by P2y12 positive microglia was unchanged in regions close to *APOE4* h-astrocytes compared with their contralateral sides (Fig 5C, D). Moreover, the area covered by Clec7a positive microglia around plaques decreased significantly in areas close to *APOE3* h-astrocytes while it increased in regions next to *APOE4* h-astrocytes compared with their contralateral sides (Fig 5E, F). These data suggest that there is a crosstalk between h-astrocytes, astrocytic hAPOE and the host microglia at the plaques. Moreover, our results indicate that *APOE3* and *APOE4* h-astrocytes and the different forms of hAPOE they secrete can indeed modulate in divergent ways microglia responses to the plaques. *APOE4* h-astrocytes and hAPOE4 prompt an increased clustering and exacerbated DAM state in microglia surrounding the plaques. In contrast, *APOE3* h-astrocytes and hAPOE3 induce a reduced clustering and more homeostatic state of microglia around plaques characterized by upregulation of the homeostatic marker P2y12 and downregulation of the DAM marker Clec7a.

### *APOE3* h-astrocytes ameliorate while *APOE4* h-astrocytes aggravate neuritic dystrophy and Tau pathology

Large swollen dystrophic neurites and hyperphosphorylated Tau are associated with Aβ plaques in AD patients and mouse models. Therefore, we wondered whether *APOE3* and *APOE4* h-astrocytes and secreted hAPOE could influence the degree of amyloid-associated neuronal toxicity and Tau pathology in chimeric AD mice. We analyzed neuritic dystrophy around Aβ plaques of similar sizes (Fig S7E) by immunostaining with the monoclonal antibody APP, which accumulates within dystrophic neurites. We measured the area of APP positive dystrophic neurites around Aβ plaques. Quantification of APP surrounding Aβ plaques in cortical areas close to h-astrocyte transplants revealed a significant decrease in the area occupied by dystrophic neurites in cortical regions hosting *APOE3* h-astrocytes compared with contralateral regions (Fig 6A, C; Fig S6A). In contrast, we found an increase in the area occupied by dystrophic neurites surrounding plaques in cortical regions close to *APOE4* h-astrocytes compared with contralateral sides (Fig 6A, C; Fig S6A).

**Figure 6.**
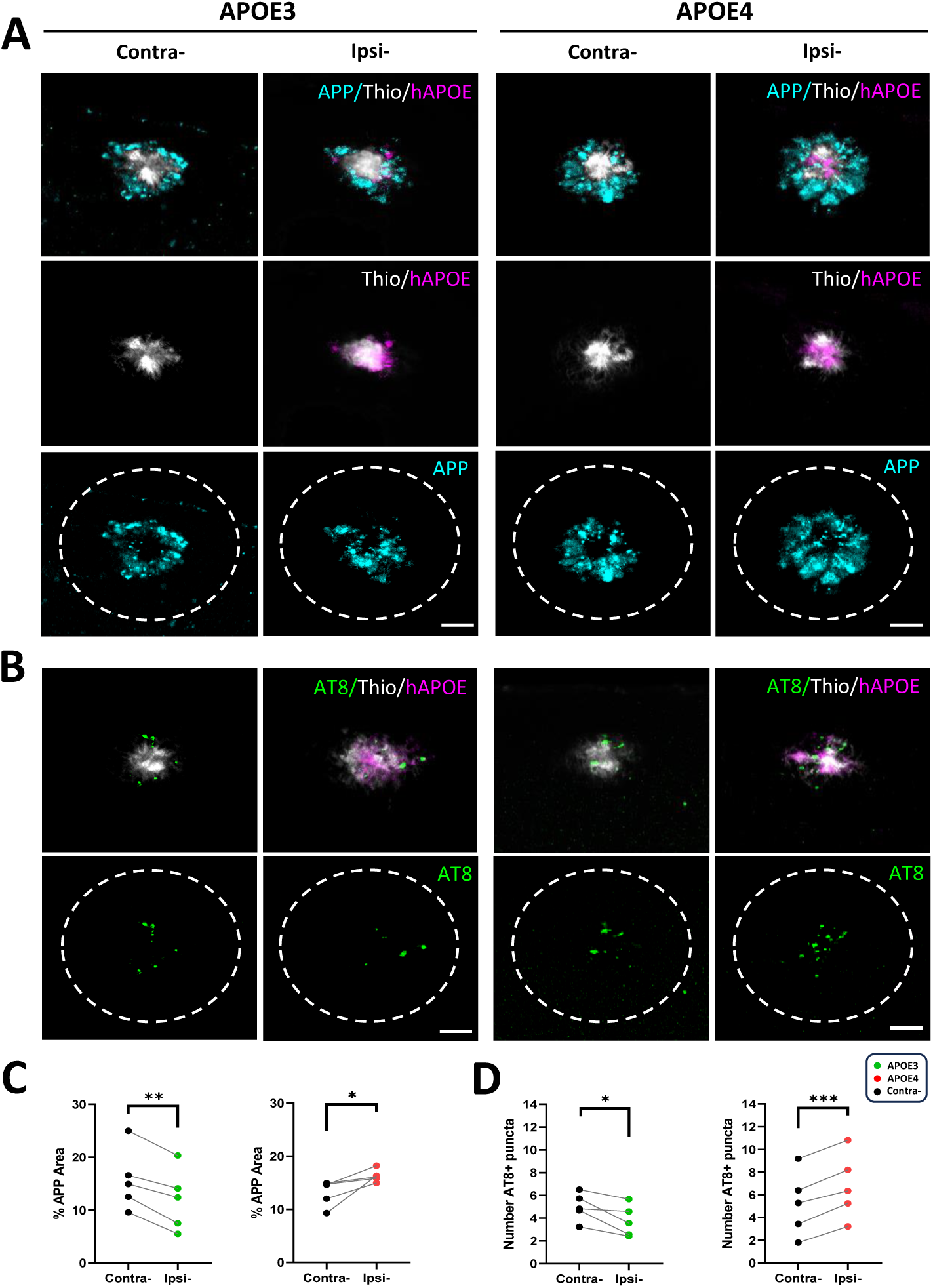
*APOE3* h-astrocytes ameliorate while *APOE4* h-astrocytes aggravate neuritic dystrophy and Tau pathology. **(A)** Representative images of dystrophic neurites (APP, cyan) surrounding Aβ plaques (Thio, white) with and without human APOE (hAPOE, magenta) in contralateral and ipsilateral cortices of AD chimeras 5-6 months after transplantation. Scale bars: 10 μm. **(B)** Representative images of hyperphosphorylated Tau (AT8, green) surrounding Aβ plaques (Thio, white) with and without human APOE (hAPOE, magenta). Scale bars: 10 μm. **(C)** Percentage area occupied by APP positive dystrophic neurites around plaques in contra- and ipsilateral cortices (n=5 mice with *APOE3* and n=5 mice with *APOE4* h-astrocytes). **(D)** Number of AT8 positive puncta around plaques in contra- and ipsilateral cortices (n=5 mice with *APOE3* and n=5 mice with *APOE4* h-astrocytes). Data are represented as mean ± SEM, paired t-test; ns = not significant, * = p < 0.05, ** = p < 0.001, *** = p < 0.0001.

Similarly, we analyzed Tau pathology around Aβ plaques in cortical areas next to h-astrocyte transplants and contralateral regions by immunostaining with the p-Tau specific monoclonal antibody AT8. Interestingly, we found a reduced number of AT8 positive puncta around hAPOE positive Aβ plaques in areas close to *APOE3* h-astrocytes compared with contralateral areas (Fig 6B, D; Fig S6B, C). However, there was an increased number of AT8 puncta around plaques close to *APOE4* h-astrocytes compared with the contralateral side (Fig 6B, D; Fig S6B, C). Overall, these results further point to a differential impact of *APOE3* and *APOE4* h-astrocytes and secreted hAPOE on Tau pathology and neuritic dystrophy around Aβ plaques in chimeric mice. While *APOE3* h-astrocytes and hAPOE3 reduce both Tau pathology and neuritic dystrophy, *APOE4* h-astrocytes and hAPOE4 promote these AD processes having a critical impact on AD progression.

In summary, our data highlight the important contribution of h-astrocytes and astrocytic hAPOE to main AD pathological hallmarks in chimeric mice and reveal a distinct impact of h-astrocytes with different *APOE* variants to main AD hallmarks and AD progression. Importantly, *APOE3* h-astrocytes and hAPOE3 ameliorate not only Aβ pathology but also other key AD processes such as Tau pathology and neuritic dystrophy. In contrast, *APOE4* h-astrocytes and hAPOE4 exacerbate these pathological features. Moreover, they modulate microglial responses to Aβ pathology in opposite ways. *APOE4* h-astrocytes and hAPOE4 enhance microglia clustering and exacerbate DAM state whereas *APOE3* h-astrocytes and hAPOE3 reduce clustering and induce a more homeostatic state on plaque-associated microglia.

## DISCUSSION

In this work, we have built further on previous efforts ^5^ and have generated chimeric mice transplanted with isogenic h-astrocytes harboring either *APOE3* or *APOE4* alleles. We have found that *APOE3* and *APOE4* h-astrocyte progenitors differentiate into mature astrocytes within the mouse brain and often integrate in upper layers of the cortex. A large proportion of both *APOE3* and *APOE4* h-astrocytes show morphological features of interlaminar astrocytes present in Layer 1 of the human cortex ^27,28^, in agreement with our previous findings ^5^. We speculate that interlaminar astrocytes, which are exclusively found in hominids^34^ and have very long processes descending into deeper cortical layers, have an important role in long-distance signaling within the cortex in physiological conditions and a critical role modulating AD progression.

Since astrocytes are major producers of APOE in the brain, we examined whether h-astrocytes were expressing and secreting hAPOE *in vivo*, in the host brain. We also investigated whether there were isoform-dependent differences in hAPOE expression and secretion between *APOE3* and *APOE4* h-astrocytes. Interestingly, we have found that *APOE4* h-astrocytes express and secrete lower levels of hAPOE than isogenic *APOE3* h-astrocytes in both wildtype and AD chimeric mice. These results point to an isoform-dependent regulation of hAPOE expression and secretion in h-astrocytes *in vivo*. Consistently, studies in both human and mouse astrocytes in culture have reported similar reductions in the expression and secretion of APOE in *APOE4* relative to *APOE3* astrocytes that could be due to post-translational modifications ^10,35^. Moreover, we have noticed that *APOE3* h-astrocytes upregulate hAPOE expression and secretion in AD compared with wildtype mice. However, hAPOE levels remain unchanged in *APOE4* h-astrocytes suggesting that such differences in expression and secretion of hAPOE by h-astrocytes could influence AD progression. Remarkably, despite such differences, similar levels of hAPOE3 and hAPOE4 are found at the plaques, indicating that hAPOE4 binds with higher affinity to Aβ plaques. Consistently, the ratio of hAPOE within the plaques versus secreted hAPOE surrounding the plaques was higher in cortical areas close to *APOE4* h-astrocytes than in areas next to *APOE3* h-astrocytes. We extended these analyses to human AD patient samples and observed hAPOE in Aβ plaques similar as in chimeric mice. The presence of APOE in the plaques in both AD mice and AD patients has also been described in a recent study ^36^ that suggests that APOE is indeed necessary for plaque formation. Moreover, a higher hAPOE ratio was also present at the plaques in the lateral temporal cortex of *APOE4* patients compared with *APOE3* AD patients, confirming a higher affinity of hAPOE4 to bind and accumulate within the plaques in AD patients as well, and highlighting the translatability of our findings.

We also aimed to answer the question of how *APOE3* and *APOE4* h-astrocytes and/or hAPOE3 and hAPOE4 specifically produced by them influence **main AD pathological hallmarks**. Importantly, we found divergent effects of *APOE3* and *APOE4* h-astrocytes on **A**β **pathology**. Cortical regions hosting *APOE3* h-astrocytes showed reduced Aβ burden as well as reduced soluble and insoluble Aβ_42_ levels. However, regions with *APOE4* h-astrocytes had more Aβ plaques per area compared with their respective contralateral sides. These data suggest that *APOE4* h-astrocytes are notably less efficient than their *APOE3* counterparts uptaking and clearing Aβ from the extracellular space in an *in vivo* context, as described in h-astrocytes in culture ^10^. While these effects could be solely due to differences in h-astrocyte functionality processing Aβ, similar results were observed in cortical areas adjacent to h-astrocytes suggesting that both h-astrocytes and hAPOE modulate Aβ pathology *in vivo*. Interestingly, we could not detect soluble or insoluble Aβ_42_ levels in wildtype chimeras. These data indicate that at this stage, neither *APOE3* nor *APOE4* h-astrocytes are producing Aβ to levels sufficient to be detected in our assays. This rules out that the Aβ from the transplanted astrocytes is a contributing source to plaque burden in our model. In addition, plaques in regions close to *APOE3* h-astrocytes were more compact while no differences in plaque compactness were observed in plaques next to *APOE4* h-astrocytes compared to their contralateral sides. These results indicate that h-astrocytes carrying different *APOE* variants and the different forms of hAPOE they secrete are influencing in different ways the physical structure of Aβ plaques ^9^. In line with these results in chimeras, analyses of Aβ burden in AD patients at Braak stages V and VI showed increased Aβ load in the lateral temporal cortex in *APOE4* compared with *APOE3* patients, as described previously ^30,37,38^.

Moreover, we found opposing effects of *APOE3* and *APOE4* h-astrocytes on **Tau pathology and neuritic dystrophy.** Our data revealed a reduced number of AT8 positive puncta and a significant decrease in the area occupied by dystrophic neurites surrounding Aβ plaques in cortical regions next to *APOE3* h-astrocytes compared with contralateral areas. In contrast, we found an increased number of AT8 puncta and an increase in the area occupied by dystrophic neurites around plaques close to *APOE4* h-astrocytes compared with contralateral sides. As these phenotypes could be affected by the size of Aβ plaques, both Tau pathology and neuritic dystrophy were analyzed in Aβ plaques of similar sizes in *APOE3* and *APOE4* chimeras and thus changes observed were not related to plaque size. Overall, these results further highlight the **critical impact of h-astrocytes on AD progression** and point to a differential impact of *APOE3* and *APOE4* h-astrocytes and their secreted hAPOE not only on Aβ pathology but on other AD key processes as well.

One of the intriguing phenotypes we found in AD chimeras was a differential response and activation state of **microglia surrounding A**β **plaques**. In cortical areas close to *APOE3* h-astrocytes microglia depicted reduced clustering around plaques and a more homeostatic state characterized by upregulation of the homeostatic marker P2y12 and downregulation of the DAM marker Clec7a. In contrast, in areas next to *APOE4* h-astrocytes, microglia increased clustering around plaques and showed increased expression of Clec7a compared to their contralateral sides. In line with our data, recent studies in AD patients have reported increased microglia clustering around plaques in *APOE4* carriers compared with non-carriers^39^. Our results support the hypothesis that there is a direct crosstalk between h-astrocytes and host microglia and that astrocytic hAPOE is involved in signaling pathways between them that regulate in a non-cell autonomous way both microglia responses and activation states. Moreover, our results reveal that h-astrocytes harboring *APOE3* or *APOE4* alleles and the hAPOE they secrete, modulate in divergent ways the responses and activation states of plaque-associated microglia. Recent data describe a cell autonomous role of *APOE4* microglia blocking the DAM response compared to *APOE3* ^32,33,40^. Remarkably, *APOE4* h-astrocytes and hAPOE4 seem to work differently and predispose microglia toward a state that interferes with proper microglia responses to AD associated pathology in our chimeric model. Indeed, even though microglia increased clustering around plaques and showed a more activated state in *APOE4* chimeras, these cells seem to be dysfunctional as they are not able to counteract Aβ pathology, as well as Tau pathology and neuritic dystrophy efficiently as in *APOE3* chimeras. Interestingly, recent studies identified a reactive microglia population defined by specific expression of inflammatory signals whose frequency increased with *APOE4* burden. This population, called terminally inflammatory microglia (TIM), is functionally impaired, shows defects in Aβ clearance and an exhausted-like state, and has been found in both murine and human AD ^41^. While further studies to dissect the states of microglia in our chimeric model are required, we already speculate that microglia in *APOE4* chimeras might be functionally impaired, and close to an exhausted-like or TIM-like state.

Taken together, we provide evidence supporting a crucial, impactful role of h-astrocytes on AD progression and neurodegeneration. Our data also validate the use of chimeric mice as a powerful technology to explore the contribution of h-astrocytes harboring different GWAS variants to AD and investigate interactions of these h-astrocytes with other brain cells.

### Limitations of the study

First, we used immune deficient mice that allow transplantation and survival of h-astrocytes within the host brain. While these mice lack the adaptive immune system, innate immunity remains largely intact in the model, as we demonstrated previously ^42,43^ and in this study. Besides, h-astrocytes are grown within the rodent brain, which is arguably a more physiological context for human cells than 2D and 3D cell culture systems which also lack adaptive immunity. Next, our analyses of AD patient material are limited due to the small cohort of patient samples we used. While various studies in bigger cohorts of AD patient samples support our findings, additional research will be needed to further confirm the phenotypes we described in this study. Lastly, the current study does not provide information about the molecular states of h-astrocytes, host microglia and other host cell types in chimeric brains. In the future it will be thus critical to perform RNA sequencing analyses at single-cell or single-nuclei resolution and/or spatial transcriptomics to dissect the cellular states of transplanted h-astrocytes and surrounding host cells at this stage of disease. Better understanding of both h-astrocytes and microglial states and the crosstalk between them in chimeric brains could lead to new treatment strategies for AD aimed at reprogramming of both populations towards states halting AD progression and neurodegeneration.

## CONCLUSION

We describe here a promising model that allows studying *APOE3* and *APOE4* h-astrocytes *in vivo* in an AD brain environment. Importantly, we reveal an unexpected critical contribution of h-astrocytes to Alzheimeŕs pathology in AD chimeric mice. Moreover, we show that h-astrocytes with different *APOE* variants and/or the APOE they secrete differentially affect, not only Aβ burden, but also other key pathological processes such as Tau pathology, neuritic dystrophy and, remarkably, microglia responses to Aβ plaques, thus having a more impactful role on AD progression and neurodegeneration than previously thought. This can potentially have important implications when developing novel astrocyte-specific targeting therapeutics.

## RESOURCE AVAILABILITY

### Lead contact

Further information and requests for resources and reagents should be directed to and will be fulfilled by the lead contact, Amaia M Arranz.

### Materials availability

Reagents generated in this study are available from the lead contact with a completed Material Transfer Agreement.

### Data and code availability

- All data reported in this paper will be shared by the lead contact upon request.
- This study did not generate any code.
- Any additional information required to reanalyze the data reported in this paper is available from the lead contact upon request.

## ACKNOWLEDGMENTS

This work was supported by the Ministerio de Ciencia e Innovación under grant no. MCIN/AEI/10.13039/501100011033 (PID2021-125443OB-100 also by FEDER, UE and RYC2020-029494-I by FSE invierte en tu futuro), the Alzheimer’s Association (AARG-21-850389), and the Basque Government (PIBA-2020-1-0030). Human iPSC astrocyte progenitor derivation was supported by NIH NIA R01AG082362, R01AG083941, and U19AG069701. Mouse experiments were supported by the animal facility of the University of the Basque Country (UPV/EHU). Confocal microscopy was performed at the Achucarro Imaging Core. We want to particularly thank the patients and the Biobank Banco de Tejidos CIEN for their collaboration.

## AUTHOR CONTRIBUTIONS

A.M.A. conceived the project. A.M.A. and J.C.S. designed experiments. A.M.A., B.D.S. and E.A. supervised the study. J.C.S., M.M.A., M.A.T., J.O., L.R., N.G.G., P.P. and A.S. performed and analyzed experiments. L.C. and L.E. helped with acquisition of confocal images. T.S. and T.C.S. provided the mice. L.R.A. and A.R.G. provided human brain samples and performed the neuropathological examination of human cases. J.TCW. and A.G. provided the hiPSC lines and astroglial progenitors. A.M.A. and J.C.S. prepared figures and wrote the paper. All authors interpreted data and approved the final version.

## METHOD DETAILS

### hiPSC Lines

Eight hiPSC lines were generated from three *APOE E4/E4* carriers diagnosed with AD (Table 1) as described previously ^5,11^. hiPSCs were maintained in mTeSR1 plus medium (STEMCELL Technologies) on 6-well plates precoated with Matrigel (Corning). Medium was changed daily, and cells were routinely passaged when 80-90% confluent using Accutase (STEMCELL Technologies) supplemented with 10 µM Thiazovivin (Tzv, Millipore). All lines were characterized for pluripotency, capability to differentiate into different CNS cell types ^11^, and G-banding karyotyping was confirmed normal ^5^.

### Generation of Reporter hiPSC-derived Astrocyte Progenitors

hiPSCs plated in Matrigel coated plates and cultured in mTeSR1 medium were differentiated into neural progenitor cells (NPCs) by dual SMAD inhibition (0.1 μM LDN193189 and 10 μM SB431542) in embryoid bodies (EB) media (DMEM/F12 (Invitrogen, 10565) with 1x N2 (Invitrogen, 17502-048) and B27-RA (Invitrogen, 12587-010)). Rosettes were selected after 14 days in vitro (DIV) using Rosette Selection Reagent (StemCell Technologies) and additionally directed into forebrain NPCs using EB media supplemented with 20 ng/ml FGF2 (Invitrogen). Magnetic-activated cell sorting (Miltenyi Biotec) was used to enrich for the NPCs (CD271−/CD133+) ^44^. NPCs were validated by immunocytochemistry using specific markers (SOX2, PAX6, FoxP2, and Nestin, Table 2). Forebrain NPCs were dissociated using Tryple Express (Gibco, 12605028), seeded at a density of 1,000,000 cells per well in 12-well plates, transfected with the lentiGuide-tdTomato plasmid (Addgene #99,376), and selected by hygromycin. Subsequently, purified fluorescent NPCs were seeded at a low density (15,000 cells/cm2) on Matrigel-coated plates and differentiated into astrocytes in astrocyte medium (ScienCell, 1801), as described previously ^23^. Astrocyte progenitors were cultured in vitro until DIV 40-44, their identity was confirmed by immunocytochemistry and/or fluorescence-activated cell sorting (FACS) for astrocyte-specific markers (Table 2), and they were used for transplantation experiments.

**Table 2.** Antibodies used in this study. The table summarizes information about supplier company, catalog number and concentration of use.

| Antibody | Company | Catalog # | Concentration |
| --- | --- | --- | --- |
| 4G8 | BioLegend | 800703 | 1/5000 |
| Amyloid beta MOAB-2 6C3 | Merck | MABN254 | 1/500 |
| APC | Millipore | OP80 | 1/200 |
| APP A4 | Millipore | MAB348 | 1/200 |
| AQP4 | Alomone Labs | AQP-004 | 1/300 |
| AT8 | Thermo Fisher Scientific | MN1020 | 1/100 |
| Dectin-1 (Clec7a) | InvivoGen | mabg-mdect-2 | 1/50 |
| FoxP2 | Abcam | Ab16046 | 1/100 |
| GFAP | Synaptic Systems | 173 004 | 1/1000 |
| GFAP | DAKO | Z033401-2 | 1/1000 |
| HLA DR + DP +DQ | Abcam | Ab7856 | 1/100 |
| Human APOE | Abcam | Ab52607 | 1/500 |
| Human GFAP | BioLegend | 837202 | 1/200 |
| Human Nuclear Antigen | Millipore | MAB1281 | 1/100 |
| IBA1 | Synaptic Systems | 234 004 | 1/500 |
| IBA1 | Synaptic Systems | 234 308 | 1/500 |
| Lycopersicon Esculentum Lectin | VectorLabs | DL-1178-1 | 1/500 |
| Nestin | Abcam | ab22035 | 1/200 |
| NeuN | Synaptic Systems | 266 004 | 1/300 |
| P2Y12 | Synaptic Systems | 476 011 | 1/1000 |
| P2Y12 | Sigma | HPA014518 | 1/1000 |
| PAX6 | Abcam | Ab5790 | 1/100 |
| RFP | Rockland | 600-401-379 | 1/1000 |
| RFP | Rockland | 200-301-379 | 1/1000 |
| S100 | Abcam | ab868 | 1/200 |
| S100b | Abcam | ab52642 | 1/500 |
| SOX2 | Cell Signaling | 3579S | 1/300 |
| Vimentin | BD Pharmingen | 550513 | 1/100 |
| Anti-Mouse 488 | Invitrogen | A21202 | 1/500 |
| Anti-Mouse 594 | Invitrogen | A21203 | 1/500 |
| Anti-Mouse 647 | Invitrogen | A31571 | 1/500 |
| Anti-Mouse biotinylated | Vector Laboratories | BA-9200 | 1/250 |
| Anti Mouse HRP | DAKO | P 0447 | 1/200 |
| Anti-Rabbit 488 | Invitrogen | A21206 | 1/500 |
| Anti-Rabbit 594 | Invitrogen | A21207 | 1/500 |
| Anti-Rabbit 647 | Invitrogen | A31573 | 1/500 |
| Anti-Rabbit biotinylated | Vector Laboratories | BA-1000 | 1/250 |
| Anti-Guinea Pig 647 | Jackson ImmunoResearch | 706-175-148 | 1/500 |

### Mice

We used two mouse models suitable for transplantations that were generated as described previously ^5,43^. Briefly, *APP PS1* tg/wt mice, which express the mutated *APP* (KM670/671NL) and *PS1* (L166P) under the control of the Thy1.2 promoter1.1 ^45^, were crossed with immunodeficient NOD-SCID mice (NOD.CB17-Prkdc^scid^) that carry a single point mutation in the *Prkdc* gene ^46^. *APP PS1* tg/wt *Prkdc*^scid/+^ mice from the F1 generation were subsequently crossbred with NOD-SCID mice to generate *APP PS1* tg/wt *Prkdc*^scid/scid^ immunodeficient mice. Resulting *APP PS1* tg/wt *Prkdc*^scid/scid^ mice were then bred with NOD-SCID mice to generate either *APP PS1* tg/wt *Prkdc*^scid/scid^ (AD mice) or *APP PS1* wt/wt *Prkdc*^scid/scid^ (WT mice), which were used for transplantations and IF analyses. In addition, homozygous *App^NL-G-F^* (*App^tm3.1Tcs^*) knock-in mice ^47^ or homozygous *App^hu/hu^*(*App^em1Bdes^*) with a humanized Aβ sequence (G676R, F681Y, R684H) ^48^ were crossed with homozygous *Rag2*^-/-^ (*Rag2*^tm1.1Cgn^) knockout mice to obtain the *App^NL-G-F^ Rag2^-/-^* genotype (AD mice) or *App^hu/hu^ Rag2^-/-^* genotype (WT mice). Homozygous colonies were maintained, and crosses were conducted within the same genotype to obtain mice used for transplantations and ELISA analyses.

Animals were housed in groups of four per cage with ad libitum access to food and water in individually ventilated cages (IVC) within a specific pathogen-free (SPF) facility, where light/dark cycle and temperature were controlled. Experiments were performed on both male and female littermates. All rodent experiments were approved by the Ethics Committee of the University of the Basque Country (UPV/EHU) and were executed in compliance with the ethical regulation for animal research.

### Transplantation of hiPSC-derived Astrocyte Progenitors

Transplantation experiments were performed on neonatal AD and WT mice at postnatal days P1-P4 following previously described protocols ^5^ with minor modifications. In brief, hiPSC-derived astrocyte progenitor cells at DIV 44 were enzymatically dissociated using Accutase (STEMCELL Technologies™, 7920), supplemented with HB-EGF (100−47, Peprotech) and Rock inhibitor (Y-27632, Cells guidance systems), and injected into the frontal cortex of AD or WT mice. Pups were cryoanesthetized, and approximately 200,000 cells were injected using Hamilton syringes into two locations within the forebrain: 1 mm posterior to Bregma, 1.5 mm bilaterally from the midline, and 1.2 mm from the pial surface in *APP PS1 Prkdc* mice; and 1 mm and 2 mm posterior to Bregma, 1.5 mm unilaterally from the midline, and 1.2 mm from the pial surface in *App Rag2-/-* mice. After transplantation, the pups were allowed to recover under a heat lamp at 37 °C and then returned to their home cages until reaching the weaning age. Grafted pups were monitored daily for a week.

### Sample isolation from Chimeric Mice

For immunofluorescence (IF) analysis, mice at 5-6 months after transplantation were euthanized with CO_2_ and immediately perfused with phosphate-buffered saline followed by a 4% paraformaldehyde solution. Subsequently, the brain was extracted, postfixed in the same fixative solution overnight for up to 48 hours and sectioned into 40 μm thick coronal sections using a Leica VT1000S vibratome. Brain sections were stored in cryoprotectant solution (30% ethylene glycol, 30% glycerol, 40% PBS) at −20 °C until use.

For protein extraction, mice at 5-6 months after transplantation were anaesthetized with 100 mg/kg ketamine (Fatro) and 10 mg/kg xylazine (Calier) and animals were euthanized using cervical dislocation. Brains were surgically removed and sliced into 1 mm thick coronal sections using a cold brain matrix. Sections were collected immediately into cold PBS and placed under a fluorescent dissection stereo microscope (Leica MZ10F). The fine dissection of the RFP-positive areas and contralateral brain regions was performed, and collected tissue was snap-frozen in liquid nitrogen. Samples were stored at −80 C until use.

### Immunofluorescence in Chimeric Mice

IF staining on grafted brains was performed as previously ^5^ using primary and secondary antibodies (Table 2). Briefly, vibratome sections were washed three times with PBS to remove the residual storing solution. Antigen retrieval was performed by microwave boiling the sections in 10 mM tri-sodium citrate buffer (VWR) with Tween 20 (0,1%) at pH 6.0. After rinsing with PBS, the free-floating sections were placed in permeabilization/blocking buffer containing 5% serum, corresponding to the host species of the secondary antibody (Donkey) in PBST (PBS with 0.20% Triton X-100 in 1xPBS) for 1 hour at room temperature. After the blocking step, primary antibodies (Table 2) diluted in the blocking solution were added to the sections and incubated overnight at 4°C. The next day, sections were washed three times for

5 minutes in PBST. Respective secondary antibodies were added, and sections were incubated for 2 hours at room temperature. If appropriate, nuclei staining was performed using a specific anti-human Nuclear Antigen antibody (hNuclei) (Table 2) or DAPI (Sigma). Aβ plaques were visualized by staining with Thioflavin-S (1326-12-1, Sigma) or X-34 (SML1954, Merk); and the antibody 6C3 was also used to visualize Aβ plaques (Table 2). For X-34 staining, brain sections were permeabilized with 0.25% Triton X-100 in PBS for half an hour, incubated in X-34 staining solution (10 μM X-34 diluted in 60% PBS (vol/vol), 40% ethanol (vol/vol), and 20 mM NaOH) for 20 minutes at room temperature, and rinsed in 40% ethanol/PBS solution and PBS containing 0.1% Triton X-100 ^31^. For Thioflavin staining, brain sections were incubated with a filtered 0.05% aqueous Thioflavin solution in 50% ethanol for 5 minutes at RT, followed by gradual rinsing with 70%, 95% ethanol and water. Subsequently, the samples were washed three times for 5 minutes in PBST and mounted onto glass slides using the Glycergel mounting media (DAKO) and allowed to dry at room temperature.

### Human Brain Samples

Brain tissue samples from the lateral temporal cortex (Brodmann areas 21 and 22) of 6 AD patients carrying the *APOE3/E3* alleles, 5 AD patients carrying the *APOE4/E4* alleles and 4 non-demented individuals were provided by the Biobank Banco de Tejidos CIEN (Table 3), and they were processed following standard operating procedures with the appropriate approval of the Ethics and Scientific Committees.

**Table 3.**
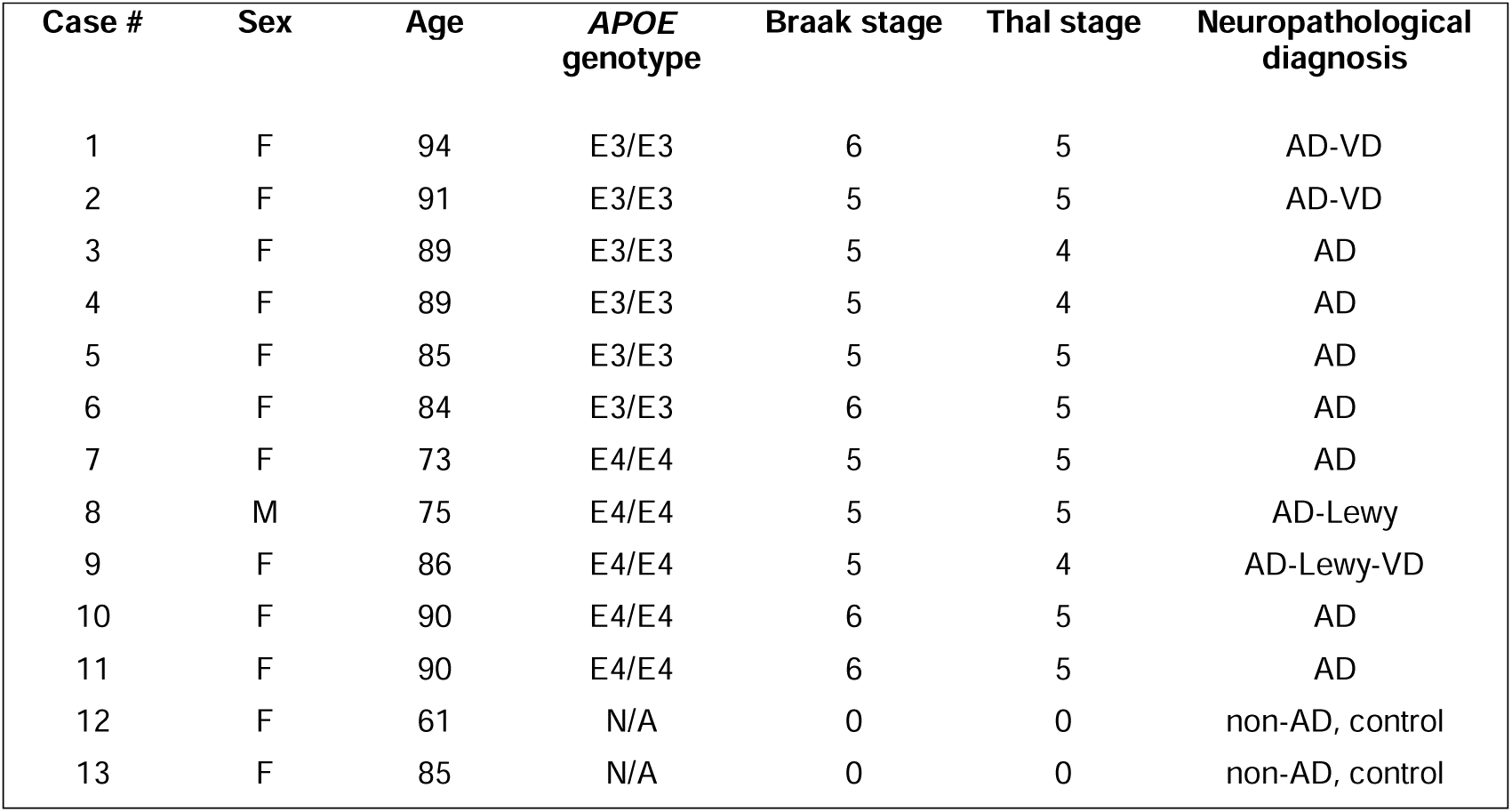
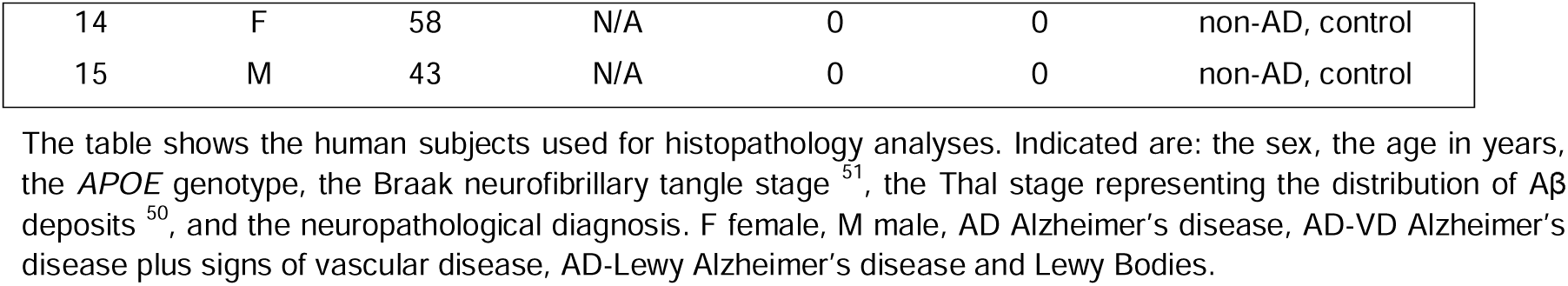
Details of Human Cases.

**Table 4.** Summary of AD-related pathology and cellular changes associated with the presence of *APOE3* and *APOE4* h-astrocytes and the hAPOE they secrete.

| **PATHOLOGY & RESPONSES** | **OBSERVATIONS IN DETAIL** | ***APOE3* vs**<br>**Contralateral** | ***APOE4* vs**<br>**Contralateral** |
| --- | --- | --- | --- |
| **Amyloid pathology** | Total Thio + plaques | ↓ | = |
|  | Total Aβ staining | ↓ | = |
|  | Aβ plaque number | ↓ | ↑ |
|  | Average plaque size | ↓ | = |
|  | Aβ plaque compactness | ↑ | = |
| **Tau pathology** | AT8 pathology around plaques | ↓ | ↑ |
| **Neuritic dystrophy** | Dystrophic neurites around plaques | ↓ | ↑ |
| **Microglia responses** | Total microgliosis | ↓ | = |
|  | Microglia clustering around plaque | ↓ | ↑ |
|  | Microglia reactivity around plaque | ↑ P2Y12<br>↓ Clec7a | = P2Y12<br>↑ Clec7a |

Brain tissues were collected as described in previous studies ^49^, with an average post-mortem interval (PMI) of 4,5 h. Briefly, after autopsy, the brains were fixed in 4% buffered formaldehyde for at least 4 weeks. Samples from the anterior entorhinal cortex and hippocampus were dissected coronally, dehydrated and embedded in paraffin. The paraffin blocks were microtomed at 5 μm, mounted on Flex IHC adhesive microscope slides (Dako), and dried at 55°C before storing. For neuropathological analysis, sections from all blocks were stained with anti-pTau (AT8), anti-Aβ (4G8) (Table 2), and with the Gallyas and the Campbell-Switzer silver techniques for detection of neurofibrillary changes and amyloid deposits ^50^. The post-mortem diagnosis of AD neuropathological change was based upon the standardized consensus criteria, including staging of Aβ plaques based on Aβ immunohistochemistry ^50^, the Braak neurofibrillary tangle (NFT) stage based on pTau immunohistochemistry ^51^, and the frequency of neuritic plaques, which compose the ABC classification system ^52^. The study comprised 15 cases with an average age of 80 years.

The table shows the human subjects used for histopathology analyses. Indicated are: the sex, the age in years, the *APOE* genotype, the Braak neurofibrillary tangle stage ^51^, the Thal stage representing the distribution of Aβ deposits ^50^, and the neuropathological diagnosis. F female, M male, AD Alzheimer’s disease, AD-VD Alzheimer’s disease plus signs of vascular disease, AD-Lewy Alzheimer’s disease and Lewy Bodies.

### Immunofluorescence in Human Brain Samples

Paraffin-embedded human sections (5 µm thick) were deparaffinized and rehydrated by immersion in xylene followed by incubations with alcohol solutions (100%, 96%, 75%, and 50% diluted in H_2_Odd) for 10 minutes each, and finally, three 5-minute washes were performed with H_2_O. Samples were then boiled in R-Universal Buffer (Aptum Biologics) using Antigen retriever 2100 (Aptum Biologics) for 20 minutes and allowed to cool down for 20 minutes to promote epitope unmasking. After retrieval, samples were washed in TBST three times for 10 minutes and blocked in 4% BSA and Tween 0,1% solution in TBS for 1 hour at RT. Following blocking, samples were incubated with primary antibodies (Table 2) diluted in the blocking solution overnight at RT. Then, samples were washed in TBST three times and incubated with fluorochrome-conjugated secondary antibodies (Table 2) diluted in the blocking solution for 1 hour at RT. Samples were washed in TBST and incubated with a filtered 0.05% aqueous Thioflavin S solution in 50% ethanol for 5 minutes at RT, followed by gradual rinsing with 70% and 95% ethanol in TBS, and water. After that, samples were washed in TBST three times and immersed in 70% ethanol for 5 minutes before being treated with Autofluorescence Eliminator Reagent (Merck, 2160) according to the manufacturer’s instructions to reduce lipofuscin-like autofluorescence. Finally, sections were washed and mounted with Fluoromount-G mounting medium (Thermofisher).

### Image Acquisition and Analyses

Confocal images were obtained using a Leica TCS STED CW SP8 confocal. For excitation, 405 nm, 488 nm, 561 nm, 638 nm laser lines were used. All the images were obtained using a 20x (0.75 NA) or 40x oil (1.4 NA) objective lens, and Z-stack series of images of the area of interest were acquired using the LAS X software. 9-10 Z-stacks with a spacing of 1 μm in chimeric brain sections and 6 Z-stacks with a spacing of 1 μm in human brain sections were obtained. All images were acquired using identical acquisition parameters such as 16-bit, 1024 x 1024 quality, and images were processed in the FIJI/ImageJ software. Images of Z-series stacks were converted to maximum intensity projections for quantifications, unless otherwise specified. Except in apoE analysis, all comparisons are between ipsilateral regions hosting h-astrocytes and their corresponding contralateral areas in the same cortical section.

#### Quantification of human APOE (hAPOE) intensity

To study hAPOE within h-astrocytes, sum projections of 40x images were used to measure mean pixel intensity per area in *APOE3* and *APOE4* h-astrocytes in WT and AD mice. To quantify the secreted hAPOE, the intensity of secreted hAPOE was measured along 100 µm from the soma of h-astrocytes, thus obtaining the intensity of secreted hAPOE relative to the distance from h-astrocyte per region. For the analysis of hAPOE in Aβ plaques, mean pixel intensity was measured in plaques within 100 µm of h-astrocytes.

#### Quantification of Aβ burden

Images were obtained with the 3DHistech Pannoramic MIDI II Slide Scanner. For quantification of the Aβ burden in regions with h-astrocytes and contralateral regions, the region with h-astrocytes and a mirror region in the contralateral hemisphere were manually selected. The % area occupied by Aβ plaques, as well as the number and size of Aβ plaques were quantified using the RenyiEntropy (for Thioflavin) or the Otus (for 6C3) algorithm in Fiji/Image J.

#### Analysis of Aβ plaque morphology

For the analysis of Aβ plaque morphology, the circularity and the Angular Second Moment (ASM) were calculated in maximum intensity projections of the z-stack images (40x). Plaques of the same size were quantified in ipsilateral and contralateral regions and a fixed threshold of 40 was used. ASM of the gray-scale co-occurrence matrix was calculated as previously ^53^ using the plugin GLCM texture of Fiji/Image J to evaluate amyloid plaque compactness. ASM serves as an indicator of local image homogeneity, with a higher value indicative of a more homogenous plaque structure with fewer transitions between light and dark areas.

#### Quantification of the plaque-associated microglia, Tau and dystrophic neurites

For quantification of the plaque-associated microglia (Iba-1, Clec7a and P2y12 signals), Tau (AT8) and dystrophic neurites (APP), images (40x) of Z-series stacks were converted to Fiji/ImageJ maximum intensity projections, and plaques with hAPOE within a maximum distance of 100 μm from the h-astrocytes were selected on each image. A 30 μm ring from the centre of mass of the plaques was drawn, except in the case of P2y12 signals in which a 40 μm ring was drawn. The area occupied by Iba-1 was quantified using the Li algorithm in Fiji/Image J. A uniform threshold, with fixed values within each animal, was then applied to measure the areas occupied by P2Y12 (20±5), Clec7a (20±5), Tau (40±5), and dystrophic neurites (40±5) within the 30 or 40 μm ring. In brief, plaques were identified with fixed threshold, and a circular area with a radius of 30 or 40 μm was generated from the center of mass of the plaque in the z-stack images. Iba-1, Clec7a, P2y12, AT8, or APP-positive area within the 30 or 40 μm ring was measured, normalized against the total expanded area around the plaque, and represented as percent positive area within the 30 or 40 μm ring. The number of AT8-positive puncta within the 30 μm ring was quantified using Analize Particles in Fiji/Image J. Plaques of the same size were quantified in ipsilateral and contralateral regions.

#### Quantification of total microglia and Aβ burden close to h-astrocytes

For the analysis of total microglia and Aβ plaque burden close to h-astrocytes, images (20x) of Z-series stacks were acquired at the periphery of the regions with h-astrocytes and converted to Fiji/ImageJ maximum intensity projections. The area occupied by microglia or Aβ plaques was quantified using a uniform threshold (35±5) with fixed values within each animal. Briefly, both plaques and microglia were identified with fixed threshold, and the positive area was measured, normalized against the total area, and represented as percent positive area.

#### Quantification of Aβ burden and APOE intensity in human cases

Individual plaques larger than 350 µm were segmented using an interactive machine learning approach, implemented through ImageJ and the Labkit plugin ^54,55^. Through this segmentation of the plaque channel, the individual plaque ROIs, the amount of Aβ, the number and size of plaques were obtained. These ROIs were also used to analyze the intensity of APOE within and around each Aβ plaque using the sum projection of the APOE channel. To measure APOE surrounding the plaques, the plaque ROI was enlarged by 10 µm, and the original plaque ROI was subtracted, leaving only the area within a 10 µm distance around the plaque.

### Protein extraction and MSD-ELISA

Protein extraction was carried out following a protocol previously described ^53^. Briefly, frozen brain tissue was homogenized in PBS with a protease inhibitor cocktail (Roche #5056489001), using FastPrep (MP Biomedicals). The homogenate was then centrifuged at 5,000 rcf for 5 minutes at 4°C and the supernatant was ultra-centrifuged at 55,000 rpm (Rotor TLA-110, Beckman Optima TLX) for 60 minutes at 4°C. The supernatant from the ultracentrifugation was used as the PBS-soluble fraction. To obtain guanidinium chloride (GuHCl) based extraction, the pellet was resuspended in 6 M GuHCl, 50 mM Tris-HCl (pHL7.6) and supplemented with cOmplete™ protease inhibitor cocktail (Roche) and sonicated using a micro-tip for 30 seconds at 10% amplitude (Branson) and then incubated for 1 hour at 25 °C. Finally, the sample was ultracentrifuged at 70,000 rpm (Rotor TLA-110, Beckman Optima TLX) and the supernatant was diluted to 0.1 M GuHCl and used as guanidinium-soluble fraction.

Aβ42 and hAPOE levels were quantified on MSD 96-well plates using MSD-ELISA (Meso Scale Discovery) as previously described ^53^. For Aβ42, standard 96-well SECTOR plates (MSD #L15XA-3) were coated with LTDA_Aβ42 capture antibody (mouse monoclonal antibody against Aβ_42_ generated in collaboration with Bart De Strooper and Maarten Dewilde labs) overnight at 4°C. The next day, the plates were washed with PBS 0.05% Tween 20 and blocked using PBS with 1% casein for 4 hours at RT. Synthetic dilutions of human Aβ_42_ were used as standards which, along with the samples, were preincubated with LTDA_hAβN labelled with a sulfo-TAG detection antibody (mouse monoclonal antibody against the N-terminal sequence of human Aβ, generated in collaboration with Bart de Strooper and Maarten Dewilde labs), in casein buffer for 5 minutes at RT. The blocked assay plate was washed five times with PBS 0.05% Tween 20, and the sample and detection antibody mix were added and incubated at 4 °C. After overnight incubation the measurements were taken. The plate was then washed with PBS, and 2x Read T buffer (MSD) was added before reading the plate on an MSD Sector Imager 2400A. For hAPOE, the R-PLEX Human APOE Assay kit (MSD #K1509MR-2) was used. Briefly, GOLD 96-well Small Spot Streptavidin SECTOR plates (MSD) were coated with biotinylated capture antibody for 1 hour at RT. After washing the plates with PBS 0.05% Tween 20, the standards and samples diluted in assay buffer were added and incubated for 1 hour at RT. Finally, the detection antibody was added and incubated for 1 hour at RT. After the incubation, the plate was rinsed and GOLD Read Buffer B (MSD) was added, and plates were read on an MSD Sector Imager 2400 A as before.

### Statistical analysis

Statistical analyses were carried out using GraphPad Prism 8 software. All data are shown as mean ± S.E.M (Standard error of the mean) with sample size and the statistical test used indicated in the figure captions. When comparing the means of two groups for a single variable, paired or unpaired two-tailed t-tests were conducted. For comparisons involving more than two groups for a single variable, a two-way ANOVA followed by Holm-Sidak’s multiple comparison test was employed. Adjusted p<0.05 (*), p<0.01 (**), p<0.001 (***) and p<0.0001 (****) were considered significant, and all significant p-values were included in Figures or noted in Figure legends.

## Supplementary Figures

**Figure S1. Transplanted cells integrate efficiently in upper layers of the cortex and express main markers of astrocytes, related to Figure 1. (A)** Representative image of h-astrocytes (RFP, yellow) expressing the human nuclei marker (hNuclei, cyan) in Layer I of the cortex of a coronal brain section of a chimeric mouse 5 months after transplantation. **(B)** Engrafted h-astrocytes (RFP, yellow) interact with blood vessels (LEL, magenta). **(C)** Engrafted h-astrocytes (RFP, yellow) express main markers of astrocytes such as GFAP (magenta), Vimentin (cyan), AQP4 (magenta), and S100β (cyan). **(D)** Engrafted h-astrocytes (RFP, yellow) show no expression of markers of oligodendrocytes (APC, magenta) or neurons (NeuN, cyan) 5 months after transplantation. Scale bars: 20 μm.

**Figure S2. Human astrocytes express and secrete APOE that binds to Aβ plaques, related to Figure 2 and Figure 3. (A)** Representative images of the chimeric mouse cortex showing *APOE3* and *APOE4* h-astrocytes (RFP, white) and human APOE (hAPOE, magenta). Scale bars: 20 μm. **(B)** Intensity profiles of hAPOE in *APOE3* and *APOE4* h-astrocytes. White lines in (A) indicate path from which the intensity profiles were generated. **(C)** MSD-ELISA quantification of hAPOE protein levels in total fractions from cortical regions with *APOE3* and *APOE4* h-astrocytes in AD mice (n=16 mice with *APOE3* and n=15 mice with *APOE4* h-astrocytes). Data are represented as mean ± SEM, unpaired t-test; ns = not significant, * = p < 0.05, ** = p < 0.01, *** = p < 0.001. **(D)** General overviews of the ipsilateral and contralateral cortical hemispheres of chimeric mice showing the presence of hAPOE (magenta) within the plaques (Thio, white) close to h-astrocytes (RFP, yellow), while no hAPOE is detected within the plaques far from h-astrocytes or in the contralateral hemisphere. Enlarged images of the inserts are shown in the bottom panel. Scale bar: 50 μm.

**Figure S3. *APOE3* and *APOE4* h-astrocytes and the hAPOE they secrete differentially affect overall Aβ burden, related to Figure 4. (A)** Representative images of the cortex of 5-6 month old AD chimeric mice stained for h-astrocytes (RFP, yellow) and fibrillar Aβ (Thio, white). Both contralateral and ipsilateral cortices hosting *APOE3* and *APOE4* h-astrocytes are shown. Scale bars: 100 μm. **(B)** Quantification of Aβ burden, number of plaques per area and plaque size in contra-and ipsilateral cortices (n=6 mice with *APOE3* and n=5 mice with *APOE4* h-astrocytes). Data are represented as mean ± SEM, Paired t-test; ns = not significant, * = p < 0.05, ** = p < 0.01, *** = p < 0.001. **(C, D)** MSD-ELISA quantification of Aβ levels in insoluble and soluble fractions from cortical regions with *APOE3* and *APOE4* h-astrocytes and contralateral areas in AD mice (C) and WT mice (D) (n=7 AD mice with *APOE3* and n=5 AD mice with *APOE4* h-astrocytes; n=10 WT mice with *APOE3* and n=8 WT mice with *APOE4* h-astrocytes). Data are represented as mean ± SEM, unpaired t-test; ns = not significant, * = p < 0.05, ** = p < 0.01, *** = p < 0.001.

**Figure S4. Human brains from *APOE4* AD patients exhibit increased Aβ load and plaque-associated hAPOE ratio, related to Figure 3 and Figure 4. (A)** Representative images of the lateral temporal cortex of AD patients carrying *APOE3* or *APOE4* alleles stained for hAPOE (magenta) and fibrillar Aβ (Thio, white). Scale bars: 50 μm. **(B, C, D)** Quantification of the Aβ plaque burden (B), the mean intensity per area of plaque-associated hAPOE (C) and the ratio of hAPOE intensity within the plaques vs hAPOE intensity surrounding the plaques (D) (n=6 AD patients carrying *APOE3/E3* alleles and n=5 AD patients with *APOE4/E4* alleles). Data are represented as mean ± SEM, unpaired t-test; ns = not significant, * = p < 0.05, ** = p < 0.01, *** = p < 0.001.

**Figure S5. h-astrocytes and secreted hAPOE modulate microglial responses in an isoform-dependent manner, related to Figure 5. (A)** Representative images of total microglia (Iba-1, green) in contralateral and ipsilateral cortices of AD chimeras hosting *APOE3* and *APOE4* h-astrocytes (RFP, yellow) 5-6 months after transplantation. Aβ plaques are stained with Thioflavin S (Thio, white). Scale bars: 100 μm. **(B)** Percentage area covered by total microglia close to h-astrocytes in AD chimeras (n=6 mice with *APOE3* and n=5 mice with *APOE4* h-astrocytes). Data are represented as mean ± SEM, paired t-test; ns = not significant, * = p < 0.05, ** = p < 0.001, *** = p < 0.0001.

**Figure S6. *APOE3* and *APOE4* h-astrocytes differentially affect overall neuritic dystrophy and Tau pathology, related to Figure 6. (A)** Representative images of dystrophic neurites (APP, red) surrounding Aβ plaques (Thio, blue) in contralateral and ipsilateral cortices of AD chimeras with *APOE3* and *APOE4* h-astrocytes. Scale bars: 25 μm. **(B)** Representative images of hyperphosphorylated Tau (AT8, green) surrounding Aβ plaques (Thio, blue). Scale bars: 50 μm. **(C)** Percentage area occupied by AT8 around plaques in contra- and ipsilateral cortices (n=5 mice with *APOE3* and n=5 mice with *APOE4* h-astrocytes). Data are represented as mean ± SEM, paired t-test; ns = not significant, * = p < 0.05, ** = p < 0.001, *** = p < 0.0001.

**Figure S7. Aβ plaque size is consistent across all analyses performed in both the ipsilateral and contralateral regions, related to Figures 4-6**. Quantification of Aβ plaque size in all analyses, comparing the ipsilateral and contralateral regions for the plaque morphology with X34 marker and the Iba-1, Clec7a, P2y12, APP and the AT8 analyses around plaques. For each analysis, Aβ plaque size in contra- and ipsilateral cortical areas with *APOE3* and *APOE4* h-astrocytes is shown, followed by the combined data for all cases. Data are represented as mean ± SEM, paired t-test; ns = not significant, * = p < 0.05, ** = p < 0.001, *** = p < 0.0001.

