## Supplementary figures and images for "Divergent Effects of *APOE3* and *APOE4* Human Astrocytes on Key Alzheimer’s Disease Hallmarks in Chimeric Mice"

### supplemental figures

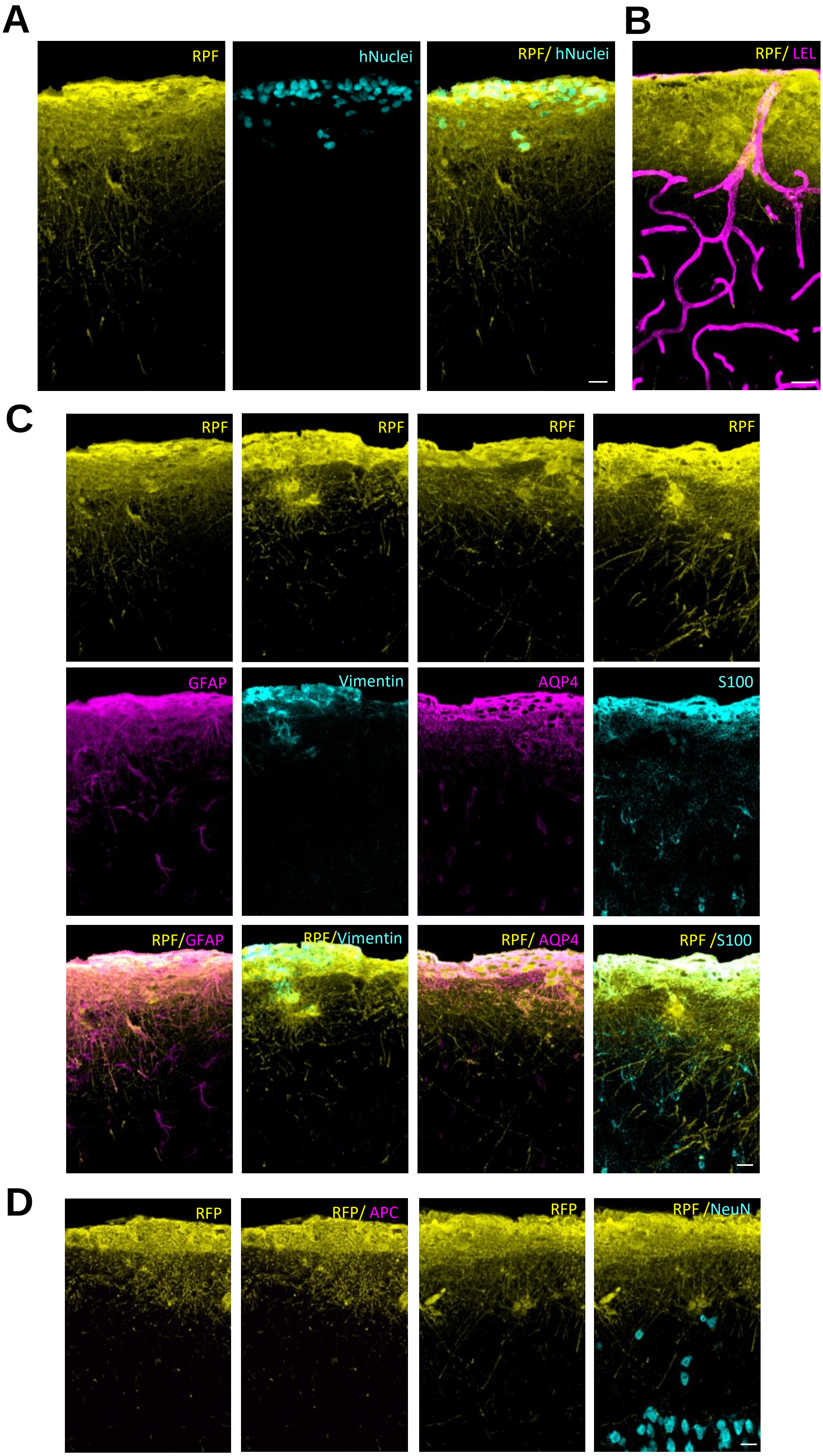

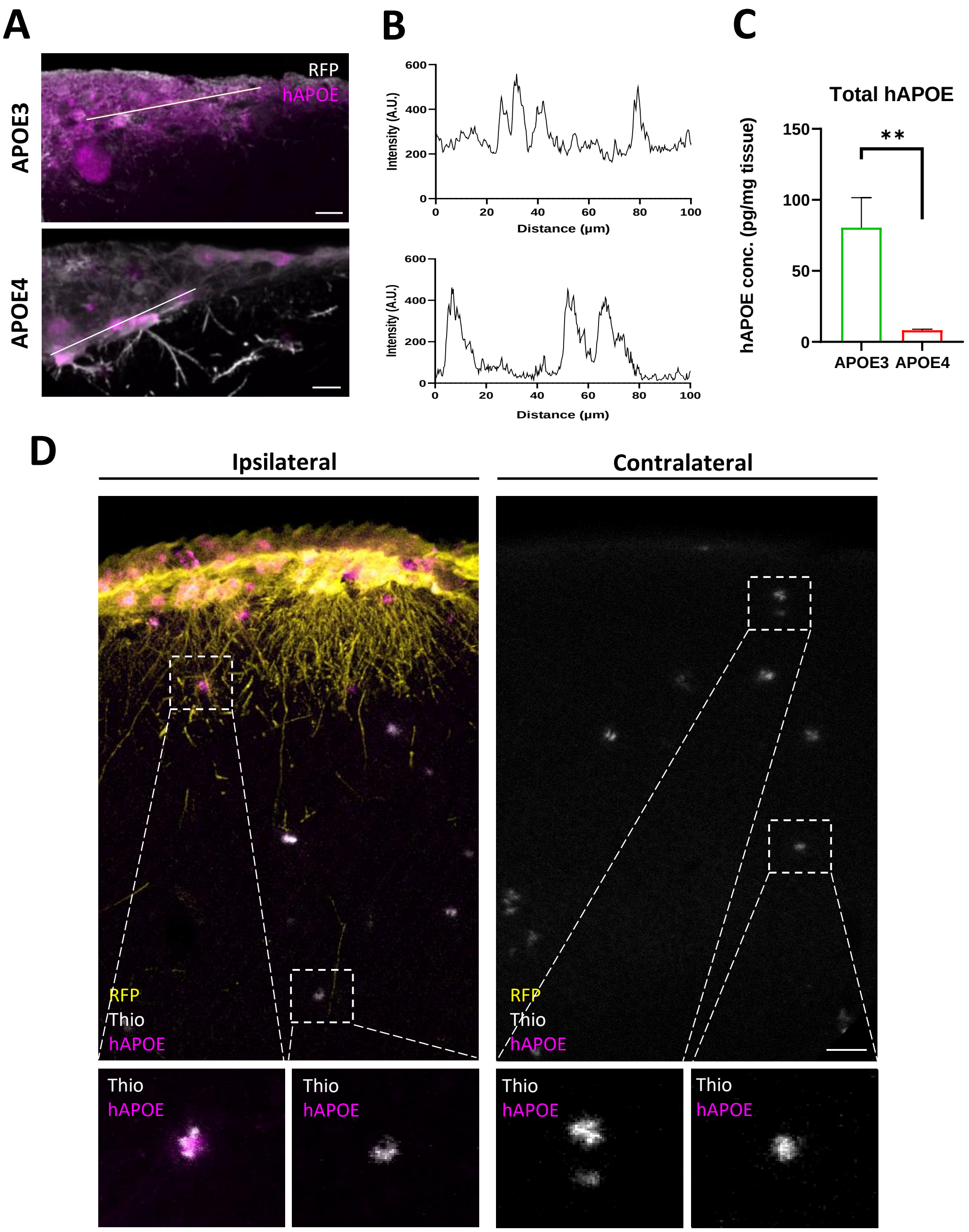

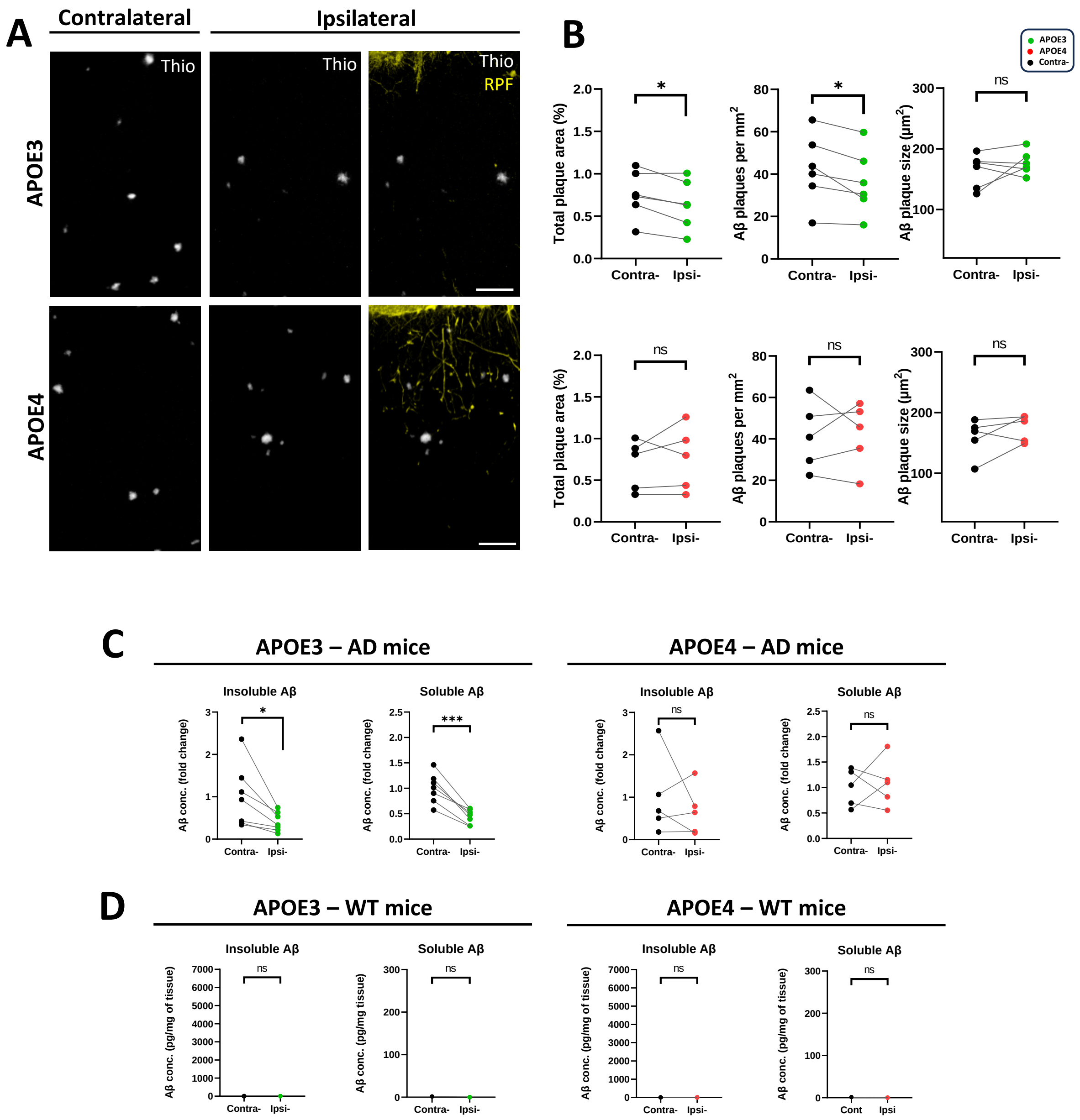

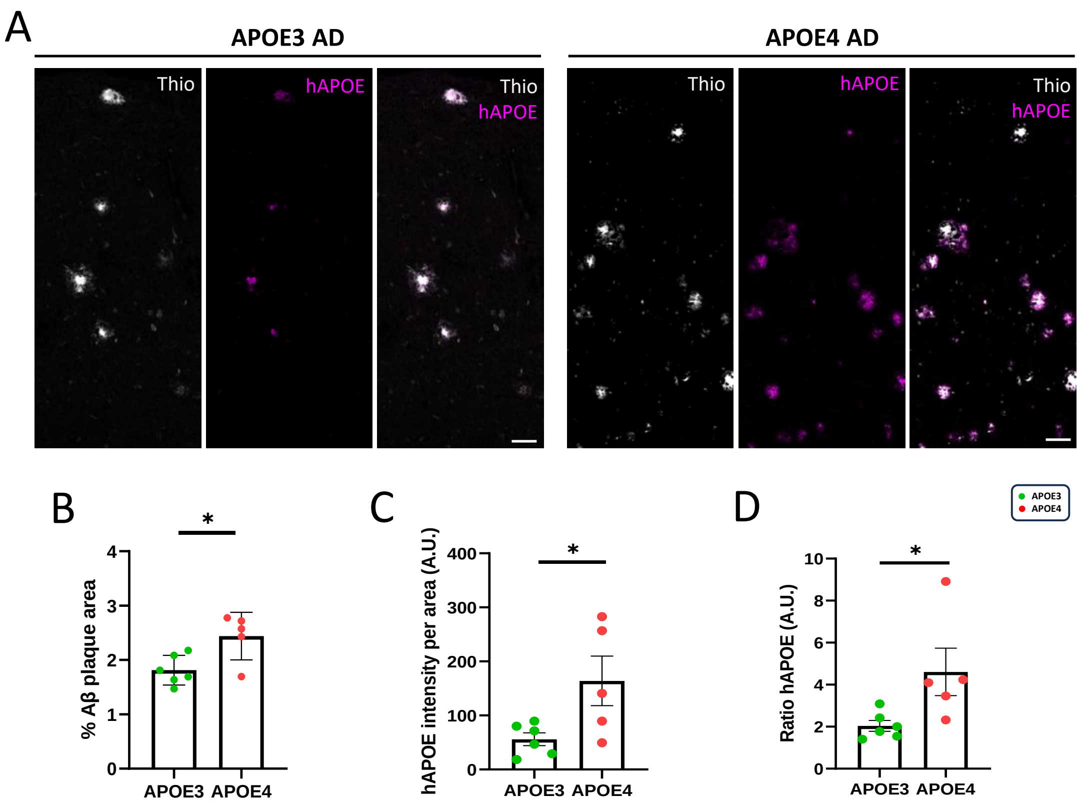

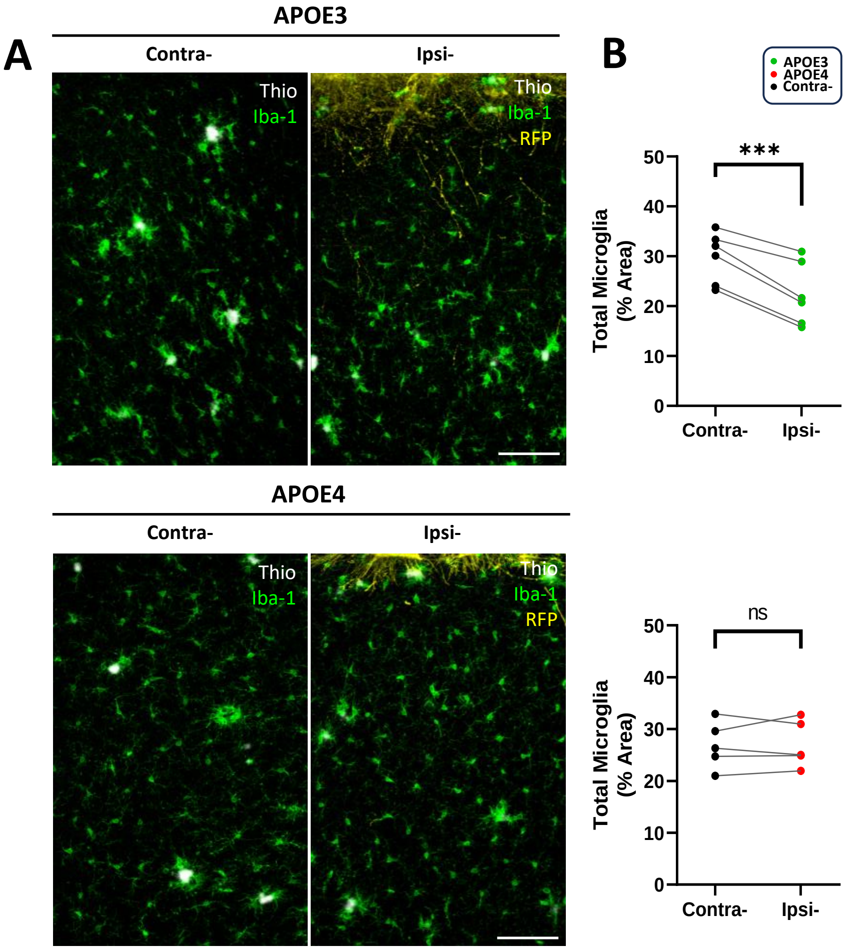

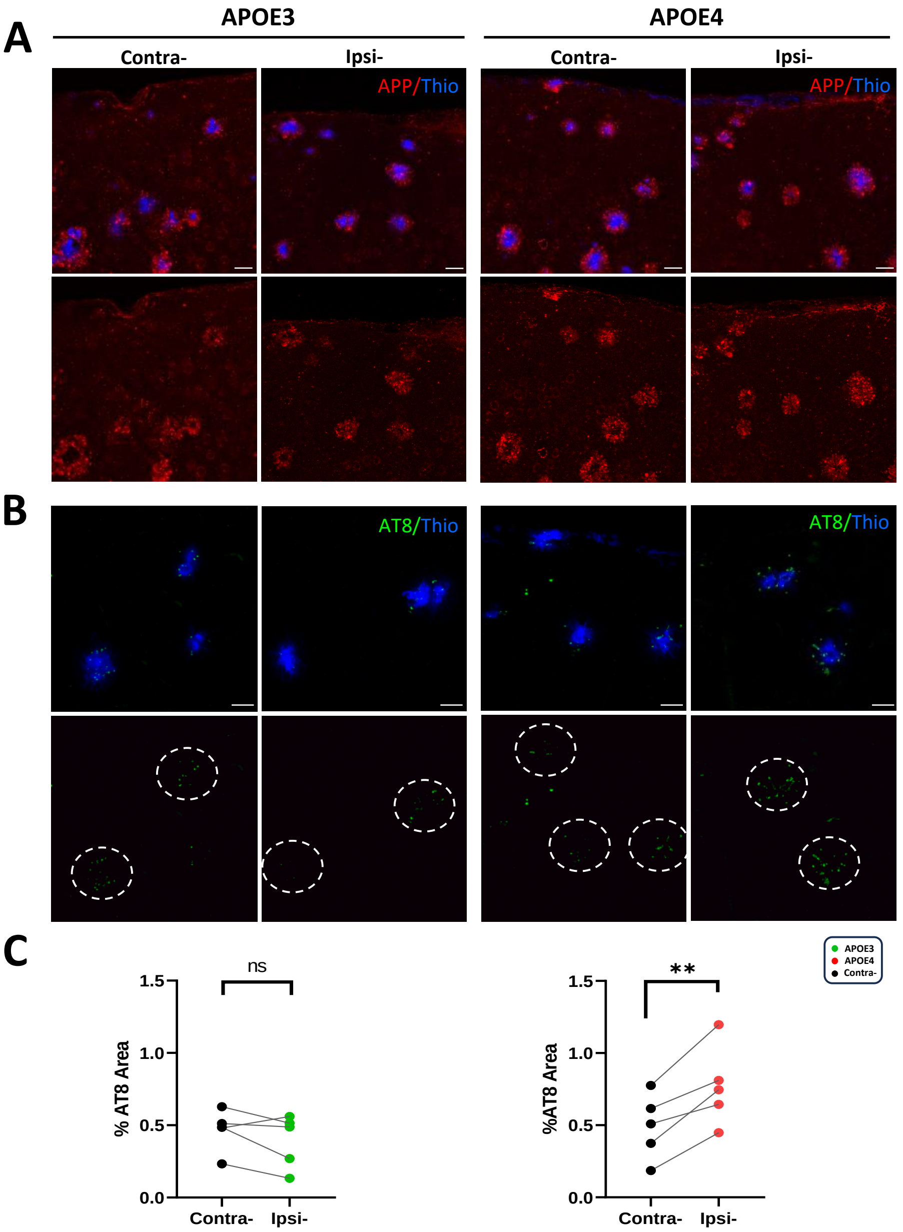

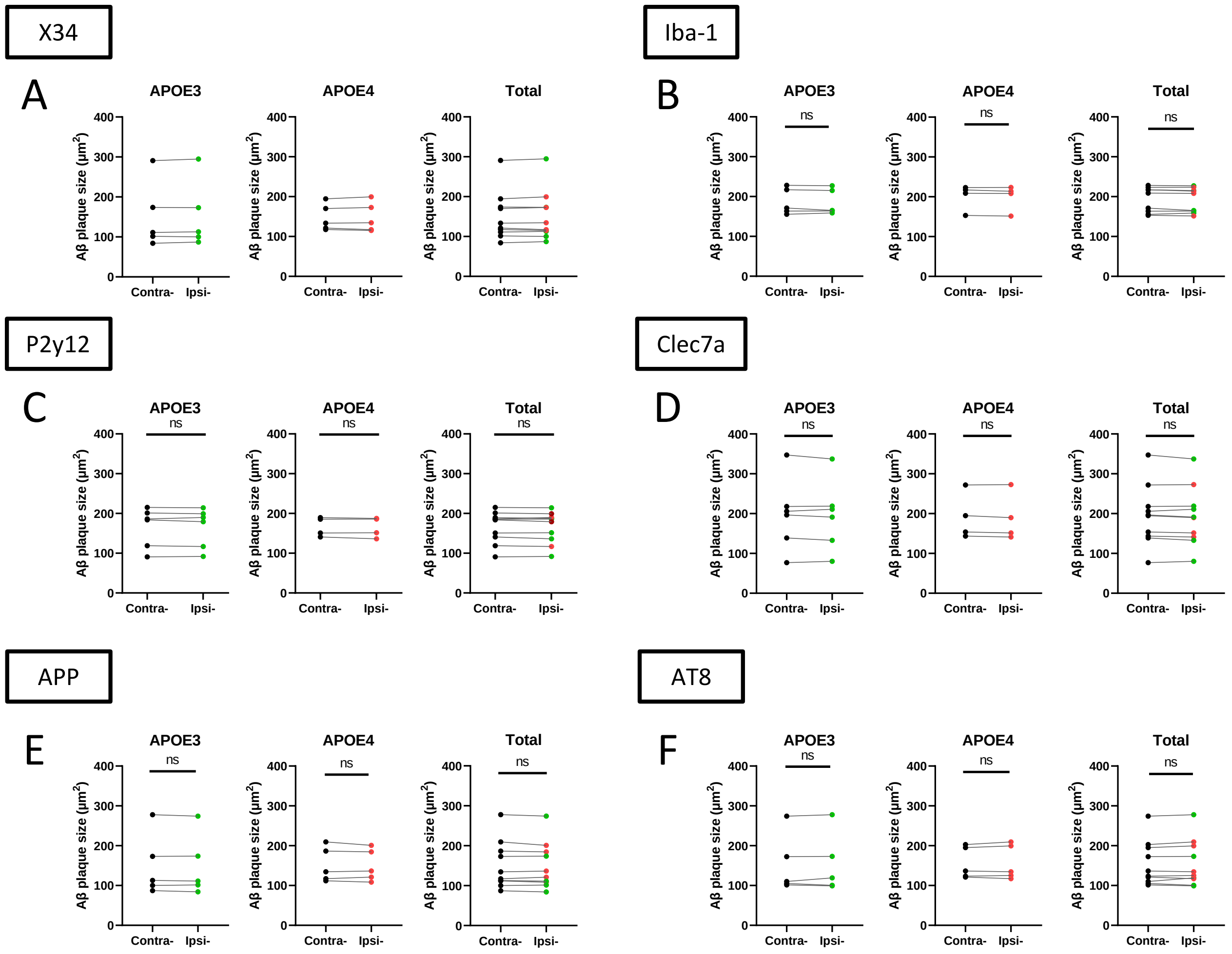
